# Salinomycin disturbs Golgi apparatus function and specifically affects cells in epithelial-to-mesenchymal transition

**DOI:** 10.1101/2022.08.31.506024

**Authors:** Marko Marjanović, Ana-Matea Mikecin Dražić, Marija Mioč, Filip Kliček, Mislav Novokmet, Gordan Lauc, Marijeta Kralj

**Author notes:** These authors contributed equally (Marko Marjanović, Ana-Matea Mikecin Dražić).

## Abstract

Epithelial-to-mesenchymal transition (EMT) gives rise to cells with properties similar to cancer stem cells (CSCs). Targeting the EMT program to selectively eliminate CSCs is a promising way to improve cancer therapy. Salinomycin (Sal), a K^+^/H^+^ ionophore, was identified as highly selective towards CSC-like cells, but its mechanism of action and selectivity remains elusive. Here we show that Sal, similarly to monensin and nigericin, disturbs the function of the Golgi apparatus (GA). Sal alters the expression of GA-related genes and leads to marked changes in GA morphology, particularly in cells that underwent EMT. Moreover, GA disturbing agents severely affect protein post-translational modifications including protein processing, glycosylation and secretion. We discover that the alterations induced by GA disturbing agents specifically affect the viability of EMT cells. Collectively, our work identifies a new vulnerability related to the EMT, suggesting that targeting the GA is a novel therapeutic approach against CSCs.

## Introduction

Epithelial-to-mesenchymal transition (EMT) is a cell transdifferentiation program during which cells lose epithelial characteristics (e.g. adhesion and lack of motility) while they acquire a mesenchymal phenotype associated with cell migration (Nieto and Cano 2012). Epithelial carcinoma cells that have passed through the EMT show properties of cancer stem cells (CSCs), which includes invasiveness, drug-resistance and the ability to form metastases at distant organs. Thereby the EMT contributes to cancer metastasis and relapse (Shibue and Weinberg 2017). Hence, targeting the EMT program to selectively eliminate CSCs is a promising way to improve cancer therapy (Pattabiraman and Weinberg 2014; Nimmakayala et al. 2019).

It was demonstrated that the induction of the EMT in immortalized human mammary epithelial cells (HMLEs) results in the expression of stem cell markers and the acquisition of the breast CSC phenotype. Consequently, these cells have been established as an experimental cell model of CSCs (Mani et al. 2008; Gupta et al. 2009). High-throughput screening of a large compound library on the HMLE cell model identified several drugs selective towards EMT/CSCs. The most potent compound was salinomycin, a potassium ionophore widely used as a coccidiostat (Gupta et al. 2009). Many subsequent studies proposed potential mechanisms of Sal’s selectivity against CSCs, as well as pursuing specific vulnerabilities of EMT and CSCs (Pattabiraman and Weinberg 2014; Kaushik et al. 2018; Antoszczak 2019; Wang et al. 2021)

Up to date, Sal has been shown to affect tumor cells through various mechanisms including activation of apoptosis, autophagy, and elevation of intracellular ROS, as well as mitochondrial membrane depolarization, and inhibition of multidrug resistance pumps. Moreover, several studies have used Sal to potentiate the efficacy of other commonly used anticancer chemotherapeutics (Huczynski 2012; Managò et al. 2015; Shi et al. 2015; Dewangan et al. 2017; Kaushik et al. 2018; Antoszczak 2019). Nevertheless, despite the enormous efforts in the past decade, the specific mechanism of Sal’s EMT/CSC selectivity remains vague. Recently, Huang et al. showed that Sal accumulates in the ER and thus promotes the release of Ca^2+^ into the cytosol. This leads to the unfolded protein response (UPR) and activation of CHOP which inhibits Wnt signaling, an effect already ascribed to Sal in CLL (Lu et al. 2011; Huang et al. 2018).

It was demonstrated that the ER-mediated UPR is permanently activated in EMT cells to support synthesis and secretion of large quantities of extracellular matrix (ECM) proteins. Hence, ER function is critical for EMT cells’ maintenance, offering a new therapeutic approach (Feng et al. 2014). In addition to increased secretory output, the EMT process involves major cellular remodeling including changes in membrane lipidome characteristics and reduced membrane fluidity, which is needed for migration and invasion (Kalluri and Weinberg 2009; Sampaio et al. 2011). Distribution of both the proteins and the lipids inside the cell is dependent on the functionally intertwined communication between the ER and the GA (Smirle et al. 2013). In addition, Mai et al. demonstrated that derivatives of Sal localize in lysosomes causing accumulation of iron and activation of ferroptosis as the main mode of CSCs cell death (Mai et al. 2017). Since Sal affects most membrane-enclosed cellular compartments, especially the ones involved in protein production, modification and secretion, as well as the fact that some ionophores with CSC selective abilities belong to the GA inhibitors (Dinter and Berger 1998; Kaushik et al. 2018), prompted us to examine the significance of the GA in EMT cells and its sensitivity to Sal. In the present study, we demonstrate that Sal affects the GA morphology as well as the expression of ER-Golgi related genes predominantly in EMT cells. Moreover, we show that Sal inhibits correct GA function which leads to altered post-translational protein modifications that are specifically processed in the GA. These include reduced secretion of proteins and marked changes in the N-glycosylation profile of secreted proteins, primarily diminishing the amount of complex N-glycans. Since our data undisputedly positions Sal as a Golgi disturbing agent and because EMT cells demonstrate greater sensitivity to the GA perturbations, we propose that targeting of the GA is a novel therapeutic path against EMT cells, and consequently against CSCs.

## Results

### EMT sensitizes cells to salinomycin and monensin

A seminal study by Gupta et al. (Gupta et al. 2009) highlighted four highly selective compounds against EMT cells: abamectin, etoposide, nigericin, and salinomycin. However, the screen yielded additional selective compounds, in particular, 32 by the algorithm used by Gupta et al. (Gupta et al. 2009). We performed a similar analysis with a more permissive threshold on the Gupta et al. (Gupta et al. 2009) data, which yielded 227 selective compounds containing several other ionophores or ionophore-like molecules (see Methods for details; Supplementary file S1A). Among these, the most interesting compound was monensin (Mon), demonstrated previously to repress the EMT gene expression signature and proposed as a potential CSCs targeting ionophore along with nigericin (Zhao et al. 2016; Kaushik et al. 2018) (Supplementary file S1B).

In the current study, we took the advantage of the same EMT models used by Gupta et al. (Gupta et al. 2009), in which non-EMT cells (HMLE-shGFP and HMLE-pBp) express epithelial marker E-cadherin and have CD24^high^/CD44^low^. Conversely, EMT cells (HMLE-shEcad and HMLE-Twist) express N-cadherin as well as CD24^low^/CD44^high^ (Fig. S1A–C). We confirmed that Sal lowered the proportion of EMT cells in a mixed population experiment (Fig. 1A and S1D). Sal also showed a more pronounced effect towards EMT cells in proliferation assays (Fig. 1B and C). The observed selectivity diminished at higher concentrations of Sal (5 μM for shEcad/shGFP or 10 μM for Twist/pBp pair) which were toxic for both EMT and non-EMT cells. Higher concentration also resulted in immediate loss of mitochondrial potential, possibly explaining the loss of selectivity (Fig. S1E). This proved its selectivity against EMT cells, although to a lesser extent than described by Gupta et al. (Gupta et al. 2009). Finally, Mon also displayed selective growth inhibition of EMT cells and enriched the percentage of cells with epithelial markers in a mixed population experiment (Fig. 1A–C, S1D). The effect was comparable to Sal. Conversely, paclitaxel (a negative control) enriched the EMT population, corroborating data from Gupta et al. (Gupta et al. 2009) (Fig. 1A and S1F). Hence, our data established that besides salinomycin and nigericin, another ionophore – monensin is an EMT selective compound.

**Figure 1.**
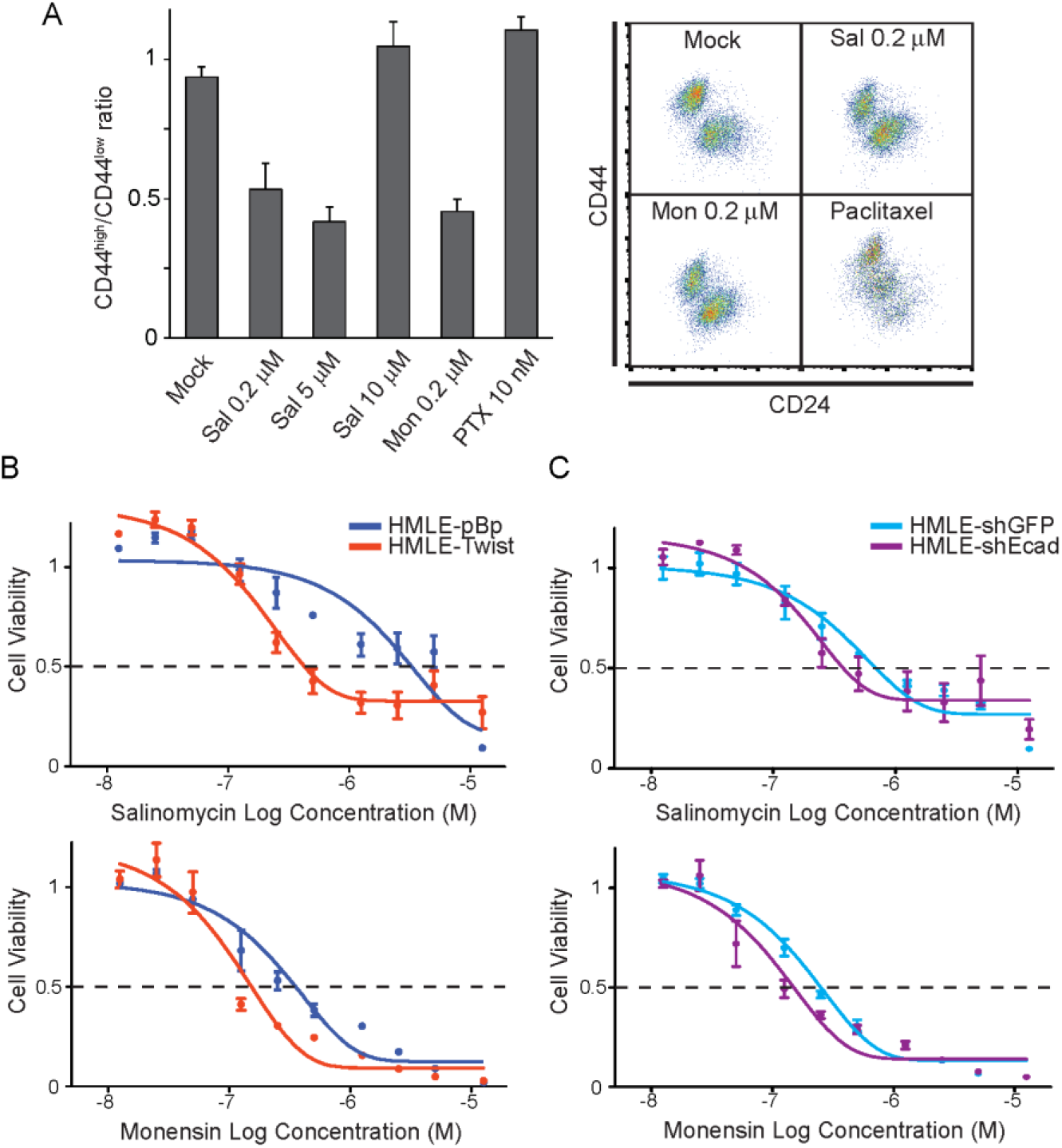
EMT sensitizes cells to salinomycin and monensin. (A) HMLE-Twist and HMLE-pBp were seeded on day 0 at 1:1 ratio and then treated with salinomycin (Sal), monensin (Mon) or paclitaxel (PTX) at indicated concentrations. Ratio of CD44^high^/CD44^low^ (EMT/non-EMT) cells after 3 days of treatment of mixed culture is shown. Representative CD24/CD44 flow cytometry profiles from are shown in the right panel. (B) Dose-response curves of HMLE-Twist (orange, EMT) and HMLE-pBp (dark blue, non-EMT) or (C) HMLE-shEcad (purple, EMT) and HMLE-shGFP (blue, non-EMT) breast cells treated with salinomycin and monensin. MTT assay was performed after 72 hours of treatment. Data are presented as mean ± s.d., n = 3 from duplicates in A and n = 3 from quadruplicates in B and C.

### Salinomycin induces the expression of ER-Golgi related genes

To evaluate the effects of Sal on intracellular compartments, we profiled the global gene expression in both EMT and non-EMT cells by RNA-Seq. We treated HMLE-Twist and HMLE-pBp cells Fig. 1A and S1D). The analysis of differentially expressed genes (≥ 2 fold change) revealed that: *i*) a common set of 52 genes were upregulated in both cell lines, *ii*) one gene (*NFE2L3*) was upregulated in EMT but downregulated in non-EMT cells, and *iii*) 3 genes (*IDI1, TM7SF2*, and *INSIG1*) were downregulated in EMT cells, but upregulated in non-EMT cells (Fig. S2A and Supplementary file S2A–D). We further widened the analysis for the case (*ii*), which yielded 64 genes (*ii+*; ≤ 0 in non-EMT and ≥ 2 fold change in EMT cells, Fig. S2B, and Supplementary file S2E). Both sets (*i* and *ii+*) include multiple genes related to the ER, the GA and membrane function, suggesting that Sal affects these cellular compartments predominantly. Observed effect can be considered to be either a general mechanism of Sal (*i*) or specific to EMT cells (*ii+*). Moreover, Gene Ontology (GO) analysis of these 2 gene sets showed enrichment of the secretory pathway-related genes (Supplementary file S2F and G, respectively).

We have additionally performed GO analysis on full sets of differentially expressed genes in the 3 experimental cases (EMT vs non-EMT, EMT with Sal and non-EMT with Sal). The analysis demonstrated that Sal treatment enriched for genes related to the ER and the GA regardless of the cell type which reinforced the notion that Sal enriches for genes involved in the secretory pathway (Fig. 2A). Additionally, a gene set enrichment analysis (GSEA) demonstrated that Sal significantly induced the expression of genes known to be upregulated by well-known GA stressors (monensin, nigericin, brefeldin A and 4MU-xyloside) or which activate the ER-related UPR (tunicamycin and thapsigargin) (Fig. 2B and C, S2C). Sal also upregulated genes reported to be enriched upon the induction of the GA-specific UPR (Serebrenik et al. 2018) (Fig. 2B and C). On the contrary, we observed no enrichment for the gene set upregulated by the treatment with the DNA damaging chemotherapeutic doxorubicin (Fig. 2B and C). In addition, Sal also induced genes involved in glycosylation, a post-translational protein modification mainly executed inside the ER-GA compartments (Fig. 2C and S2C).

**Figure 2.**
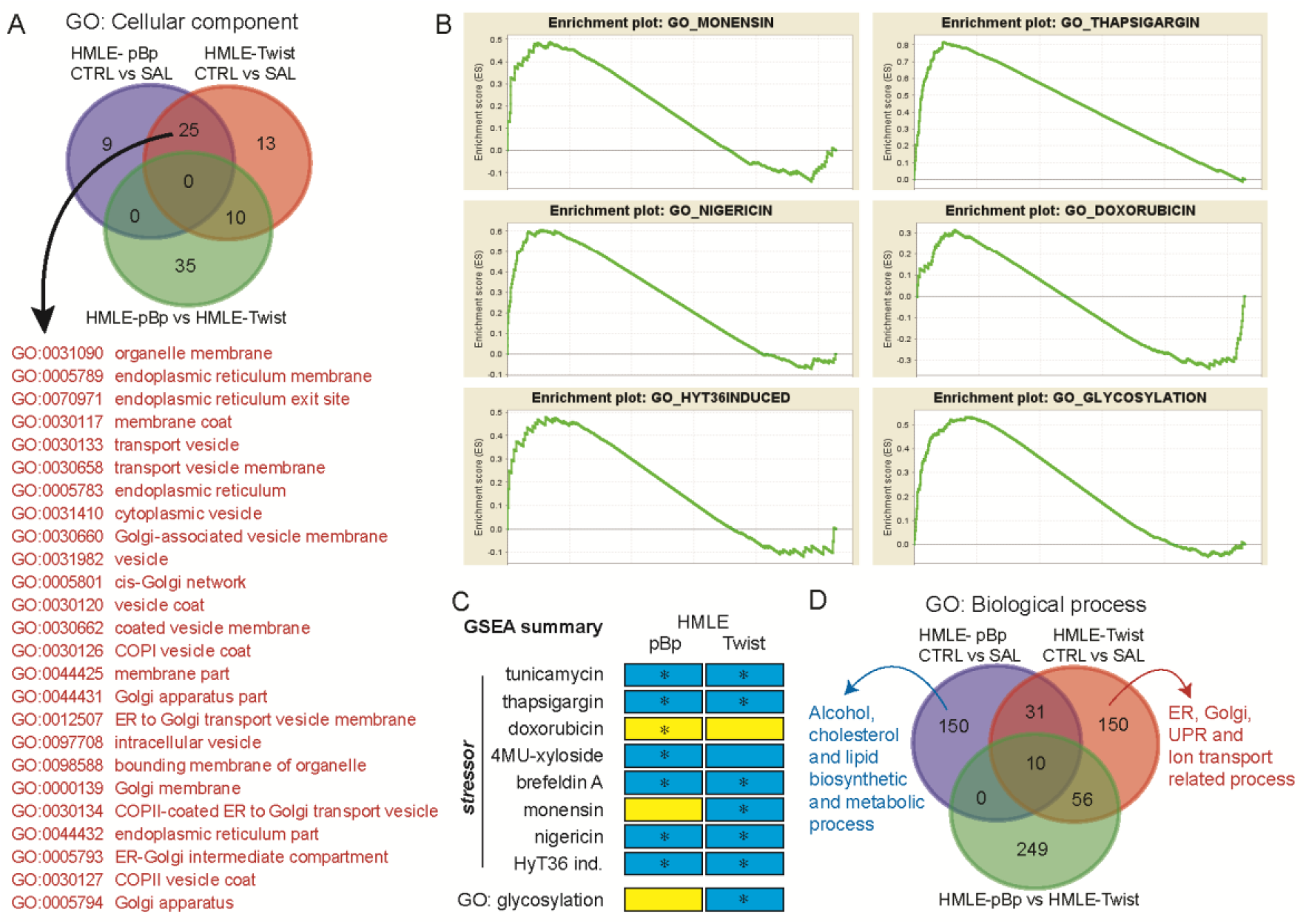
Salinomycin induces the expression of ER-Golgi related genes. RNA-seq analysis was performed on HMLE-pBp and HMLE-Twist cells (non-EMT and EMT) treated with or without 0.2 μM Sal for 24 hours in biological triplicates. Differentially expressed genes were sorted from highest to lowest in each of 3 experiments indicated and the list was analyzed with GOrilla algorithm (http://cbl-gorilla.cs.technion.ac.il/) for enrichment of cellular component (A) and biological process (D) GO categories. Names and codes for cellular component GO categories enriched in both HMLE-Twist and HMLE-pBp cells after treatment with Sal are indicated in red (A). (B) GSEA analysis for gene sets of known Golgi stressors monensin and nigericin, induction of unfolded protein response in Golgi (HyT36), ER stressor thapsigargin, DNA damage compound doxorubicin and GO category glycosylation on RNA-Seq data from HMLE-Twist cells treated with Sal is presented (analysis was performed on data ordered from largest to smallest) (see Methods for details about gene sets used). (C) Summary of GSEA analysis on different gene sets (ER and GA stressors or GO category) and their relation to enrichment in our experimental model (see Methods for details). * - significantly enriched; yellow - enrichment in mock treated sample, blue - enrichment in Sal treated sample. (D) Venn diagram for overlapping categories is shown with number indicating different GO categories. Biological processes GOs that were most enriched after Sal treatment in HMLE-Twist and HMLE-pBp cells are indicated in red and blue, respectively; full list is presented in Supplementary file S2).

Importantly, the analysis also revealed the difference between EMT and non-EMT cells in response to Sal. While GO enrichment analysis of EMT cells showed enrichment of processes related to the ER, GA and ion transport, the GO categories with the highest enrichment in non-EMT cells were related to the metabolism of alcohol, cholesterol, and lipids (Fig. 2D and Supplementary file S2H–J). Moreover, because genes upregulated in non-EMT cells and downregulated in EMT cells (set *iii*) are either directly involved in (*IDI1* and *TM7SF2*) or regulate (*INSIG1*) cholesterol biosynthesis, we assessed the global effect of Sal and Mon on cellular lipid content. We stained cells with Nile Red, a dye which can differentiate between neutral lipids (cholesterol ester or triglycerides) found in lipid droplets (LDs) (Diaz et al. 2008). We found that both compounds increased green to yellow fluorescence, representing LDs with high cholesterol content, exclusively in non-EMT cells, agreeing with the gene expression data. On the other hand, there was no obvious change observed in EMT cells (Fig. S2D). In addition, EMT cells were reported to maintain low cholesterol to allow increased membrane fluidity necessary for migration and invasion (Sampaio et al. 2011). ABCA1, a cholesterol efflux transporter, is highly expressed in EMT cells and it was demonstrated that can be downregulated by anti-metastasis drugs including salinomycin (Zhao et al. 2016). Our RNA-Seq data confirmed increased expression of *ABCA1* in EMT cells (3.45 fold change) and Sal did decrease its expression in both cell lines (Supplementary file S2A–C). Taken together, observed data suggests that Sal targets cellular compartments involved in protein and lipid processing and secretion (the ER and the GA) which activate different gene responses in EMT and non-EMT cells.

### Salinomycin affects Golgi apparatus morphology and leads to the activation of PERK branch of UPR

Next, we decided to investigate the effect that Sal has on the secretory pathway, namely the GA and the ER, as well as the induction of UPR. We inspected the GA morphology by confocal microscopy, using common GA markers GM130 and GIANTIN that co-localize with cis-Golgi and cis- to medial-Golgi matrix membranes, respectively. Sal induced fragmentation of Golgi cisternae, which appeared as a dispersed area within cells that stained positive for both GM130 and GIANTIN (Fig. 3A and B). The extent of the GA perturbations in EMT cells was statistically significant when compared to mock-treated cells (Fig. 3C and D). On the other hand, the GA of non-EMT cells did not display perturbations in morphology after Sal treatment. Of note, the dispersal of Golgi cisternae in untreated non-EMT cells was on average much larger than in EMT cells (p < 0.001 for both markers, unpaired two-tailed T-test), but the variance within the populations was very large (Fig. 3C and D). More prominent GA fragmentation in EMT cells was also observed when cells were treated with Mon (Fig. 3E).

**Figure 3.**
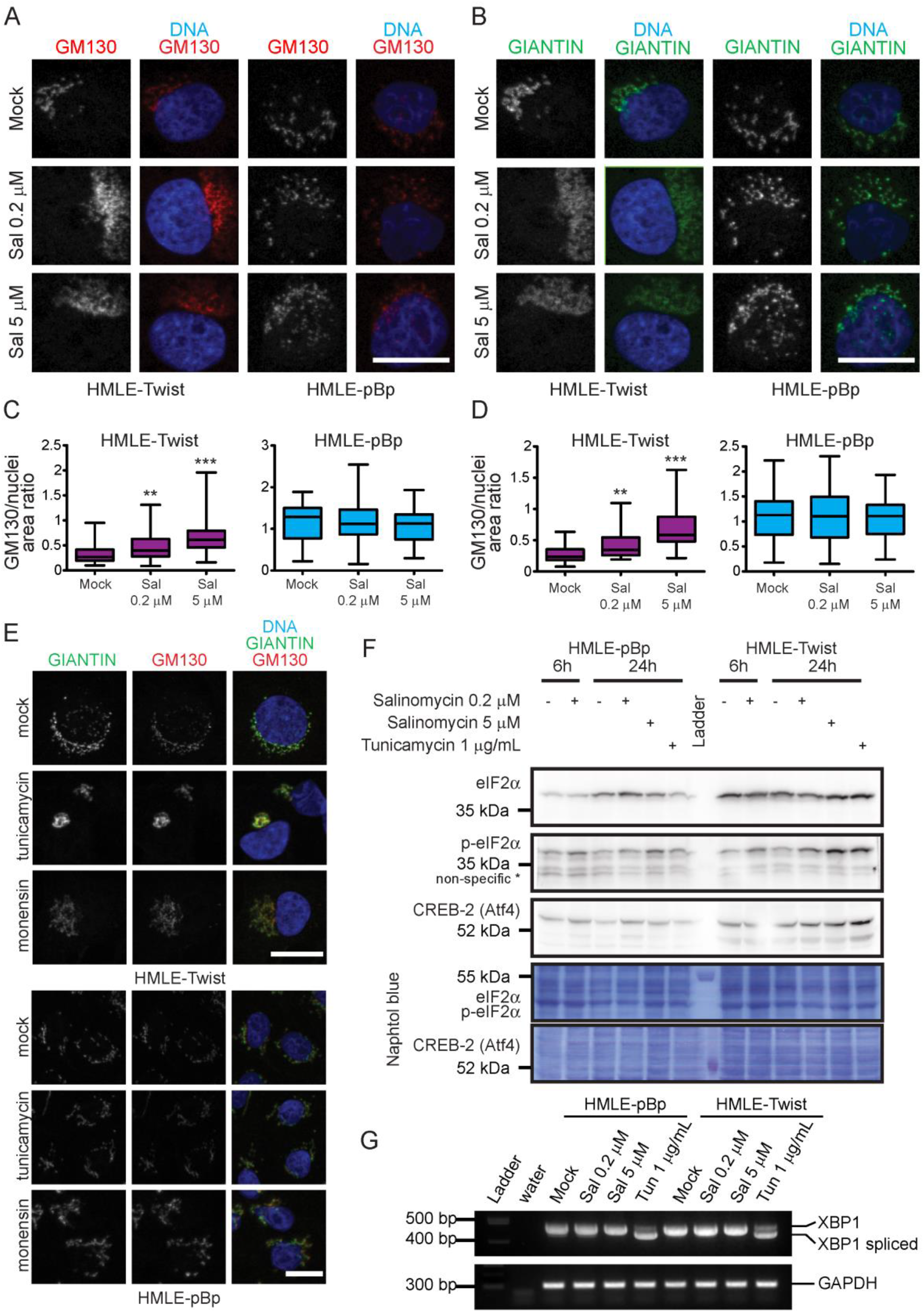
Salinomycin affects Golgi apparatus morphology and leads to the activation of PERK branch of UPR. Representative confocal microscopy images of HMLE-Twist and HMLE-pBp cells after treatment with Sal for 24 hours. Cells were stained with anti-GM130 primary and secondary Alexa Fluor 568 antibodies (red, A) or anti-GIANTIN primary and secondary Alexa Fluor 488 antibodies (green, B). DNA was counterstained with DAPI (blue). Quantification of the total area of GM130 (C) or GIANTIN (D) positive organelle and the area of the nucleus of each cell was divided and plotted. Box plot charts plotted from n = 57, 52, 72, 61, 71 and 61 cells (from left to right) from 2 individual experiments. (E) Representative confocal microscopy images of HMLE-Twist and HMLE-pBp cells after treatment with tunicamycin (5 μg/mL) or monensin (5 μM) for 24 hours. Cells were stained as in A and B. (F) Western blot analysis of UPR related proteins: phosphorylated-eIF2α (p-eIF2α), total eIF2α, and CREB-2 (Atf4) proteins in HMLE-Twist and HMLE-pBp after treatment with 0.2 μM Sal for 24 hours. Tunicamycin was used as a positive control. Naphtol blue staining of transferred proteins was used as to evaluate equal loading. (G) RT-PCR analysis of UPR-related splicing of XBP1 in HMLE-Twist and HMLE-pBp after treatment with Sal for 24 hours. Tunicamycin was used as a positive control. GAPDH was used as a loading control. Maximum projection of 10 slices is displayed. Scale bar = 10 μm. One-way ANOVA with post-hoc Dunnett’s multiple comparison test was used for statistical analysis (**p < 0.01, ***p < 0.001).

We also examined the status of the compartments between the ER and the GA, by staining the cells with antibodies for ERGIC53 and β’COP, markers of ER-Golgi intermediate compartment (ERGIC) and COPI retrograde vesicles, respectively. The microscopy confirmed that these compartments were also affected by Sal treatment and that EMT cells showed greater alterations, which were more evident on COPI vesicles (Fig. S3A).

Moreover, we also inspected Golgi morphology after treatment with tunicamycin, an early blocker of N-glycosylation that is a well-known inducer of the ER stress and UPR, and was shown to be selective against EMT cells (Feng et al. 2014). Tunicamycin led to a severe GA compaction only in EMT cells while its effect on GA morphology in non-EMT cells was negligible (Fig. 3E). Nevertheless, it induced UPR in both cell lines (Fig. 3F and G). It was previously shown that Sal can induce ER-stress and activate UPR (Yoon et al. 2013; Xipell et al. 2016), which we confirmed in our cell model. We observed the activation of PERK but not IRE1 axis of UPR signaling cascade (Fig. 3F and G). In addition, a recent publication showed that a fluorescent conjugate of Sal accumulates in the ER and leads to a massive release of the ER-stored Ca^2+^ into the cytosol (∼600% increase), which was proposed as its mechanism of action (Huang et al. 2018). This prompted us to examine changes in cytosolic calcium in our experimental model. In contrast, we observed minimal or no increase of cytosolic calcium after treatment with both Sal and Mon (5 and 0.2 μM, respectively) in our EMT model (Fig. S3B–D). Even thapsigargin, an inhibitor of sarco/endoplasmic reticulum Ca^2+^ ATPase (SERCA) that leads to depletion of the ER calcium storage, led to an increase of only 60%, hinting that HMLE cells might not be prone to such extreme Ca^2+^ releases. Collectively, these findings indicated that Sal predominantly induced severe GA stress, especially in EMT cells, which were more susceptible to the drug as compared to non-EMT cells.

### Salinomycin affects post-translational modifications and sorting of proteins

The Golgi apparatus serves as the central component of the secretory pathway, and it is the main site for the modification and sorting of the membrane and secreted proteins (Morré and Mollenhauer 2009). Thus, we investigated the fate of membrane proteins after inducing the GA stress with Sal. Classical cadherins (e.g. E- and N-) have prodomain sequences (∼130 a.a.) that guard the adhesive domains against interaction before they are activated by proteolytic processing. These sequences are cleaved off by an endoprotease in the late Golgi (Koch et al. 2004). Sal, as well as Mon and nigericin, led to the appearance of an additional band with a higher molecular weight of the EMT marker N-cadherin in western blot analysis. The effect was more prominent with a lower concentration of Sal (0.2 μM, Fig. 4A and S4A) and was observed from 24h through 72h after treatment (Fig. S4B). The same effect was observed in breast tumor cell lines with high proportion of CSC-like cells (SUM 159; Fig. 4B). Conversely, paclitaxel did not disturb proteolytic cleavage of N-cadherin corroborating the hypothesis that the consequence of Sal treatment is a GA stress-related effect. We confirmed incomplete proteolytic processing by using an N-cadherin prodomain specific antibody (Fig. 4C). Strikingly, although E-cadherin maturation also requires removal of its prodomain, the treatment with ionophores did not lead to an appearance of an additional protein band in non-EMT cells (Fig. 4A and B, S4A and B). Additionally, the E-cadherin expression increased after the treatment, especially with 5 μM Sal or 0.2 μM Mon (Fig. 4A and B).

**Figure 4.**
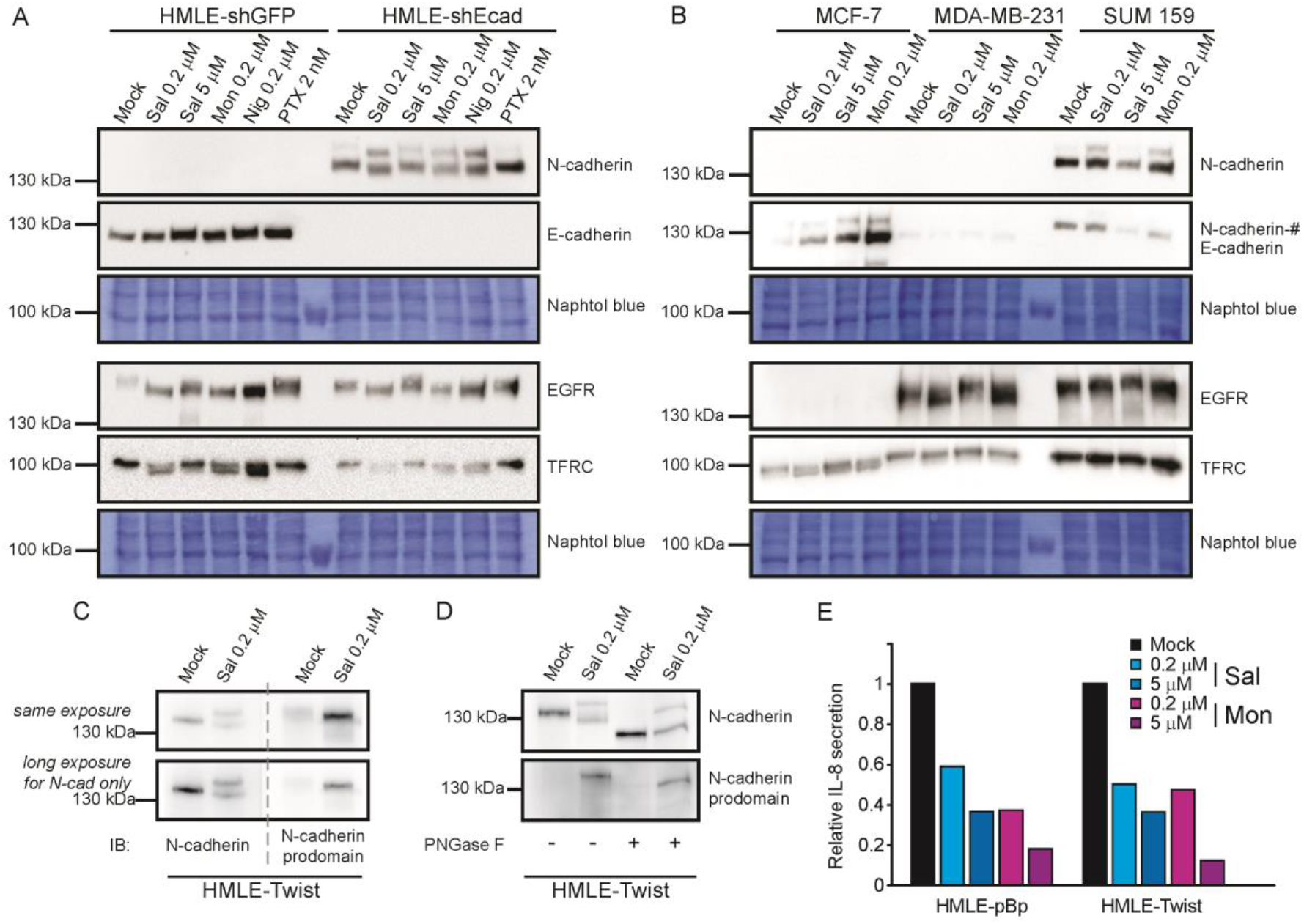
Salinomycin affects post-translational modifications and secretion of proteins. (A) Western blot analysis of membrane proteins N- and E-cadherin; epidermal growth factor receptor and transferrin receptor (EGFR and TFRC) in HMLE-shEcad and HMLE-shGFP breast cells (A) or three different breast tumor cell lines (B) that were treated for 24 hours with indicated concentrations of DMSO (Mock), salinomycin (Sal), nigericin (Nig), monensin (Mon) or paclitaxel (PTX). Naphtol blue staining of transferred proteins was used to evaluate equal loading; bloting of E-cadherin in (B) was performed after N-cadherin on the same membrane and it is showing in SUM159 cells, protein bands denoted by #. (C) Western blot analysis of proN-cadherin with specific N-cadherin prodomain antibody in HMLE-Twist cells after treatment with Sal for 24h. (D) Western blot analysis of proN- and N-cadherin in HMLE-Twist cells after treatment with PNGase F. (E) HMLE-Twist and HMLE-pBp cells were treated with DMSO (mock), salinomycin (Sal) or monensin (Mon) for 24 hours and media above cells was subjected to ELISA analysis of secreted IL-8. Data was normalized to mock-treated sample.

In addition to altered processing, we observed higher mobility of membrane proteins N-cadherin, EGFR (epidermal growth factor receptor) and TFRC (transferrin receptor) in all treated cell lines (Fig. 4A and B). We hypothesized that Sal also alters glycosylation as the ER and the GA are the major sites of protein glycosylation, a most abundant post-translational modification of membrane and secreted proteins (Varki et al. 2009). Hence, to examine the effect of reduced glycosylation of N-cadherin, we treated protein lysates with PNGase F that removes glycans attached to proteins at Asn residues (N-glycosylation). Since PNGase F removes all N-linked sugars, mobility of N-cadherin was even greater than after Sal treatment, suggesting that Sal hindered N-glycosylation (Fig. 4D).

We also examined the fate of E- and N-cadherins after treatment with a competitive inhibitor of O-glycosylation (1-benzyl-2-acetamido-2-deoxy-(α)-D-galactopyranoside (GalNAc(α)-O-bn), a known GA stressor (Miyata et al. 2013), alone or in combination with Sal. As expected, treatment with GalNAc(α)-O-bn prevented O-linked glycosylation of N-cadherin and therefore decreased the molecular mass of the mature protein (Fig S4C). Exposure to Sal and GalNAc(α)-O-bn resulted in both modifications and the mobility of both the N-cadherin prodomain and the mature protein was even greater than after treatment with Sal. On the other hand, neither the mobility nor the processing of E-cadherin was affected in non-EMT cells by GalNAc(α)-O-bn. However, the combination treatment reduced the expression and decreased the molecular mass of the mature protein to some extent. It is also noteworthy that treatment with GalNAc(α)-O-bn in combination with Sal or Mon additionally decreased the proportion of EMT cells in a mixed population experiment compared with Sal alone and showed a more pronounced effect on EMT cells in proliferation assays, whereas treatment with GalNAc(α)-O-bn alone had no effect (Fig. S4D).

In addition, we tested the influence of another compound known to disrupt Golgi apparatus function and organization by inhibiting protein transport from ER to GA, brefeldin A, together with the ER stressors tunicamycin and thapsigargin on N-cadherin and E-cadherin processing (in EMT and non-EMT cells, respectively) (Fig. S4D). Not surprisingly, tunicamycin inhibited N-linked glycosylation and therefore decreased the molecular mass of mature proteins in both EMT (N-cadherin in HMLE-Twist) and non-EMT cells (E-cadherin in HMLE-pBP and MCF -7 cells). Thapsigargin led to the appearance of an additional band with a higher molecular mass, indicating inhibition of prodomain processing in both non-EMT and EMT cells. On the other hand, surprisingly, brefeldin A also resulted in an additional band, but only in non-EMT cells, i.e., the prodomain of E-cadherin, whereas it inhibited only the glycosylation of N-cadherin, which was observed as a protein with a smaller molecular mass in EMT cells. A similar effect of thapsigargin and brefeldin A in MCF -7 cells has been described previously (Geng et al. 2012), but their different effects in EMT cells have not been described yet. These results suggest different consequences of the same treatment modality in different cell models.

We also examined the effect of the GA stressors on protein secretion. We have specifically measured the amount of interleukin-8 (IL-8), a cytokine secreted by both cell types (non-EMT cells secrete higher amount of IL-8 than EMT cells (Low-Marchelli et al. 2013)). Both Sal and Mon decreased the secretion of IL-8 by 40% or more in both cell lines (Fig. 4E). Therefore, our experiments showed that Sal hampered the post-translational modification of proteins, glycosylation and protein secretion, matching the effect observed with Mon, thus confirming Sal as a GA stressor.

### Salinomycin and monensin suppress synthesis of complex N-glycan structures

We decided to examine the N-glycome of secreted proteins for two reasons. First, we hypothesized that Sal also functions as a glycosylation inhibitor considering the observed effects on the GA, similarly to Mon, an established inhibitor of glycosylation (Dinter and Berger 1998). Second, as EMT cells were shown to depend profoundly on the increased protein secretion (Feng et al. 2014), we also hypothesized that not only diminished secretion but also a change in glycosylation of secreted proteins negatively affects EMT cell maintenance and thus represents the underlying mechanism of selectivity of these compounds. Analysis of the secretome, in comparison to the complete proteome, allowed us to overcome possible masking by uncompleted glycan structures of proteins undergoing the process of glycosylation in the cellular compartments.

The major glycan structural features in EMT and non-EMT cells were mostly similar, with both having a similar distribution of N-glycan types (oligomannose, hybrid and complex), similar relative amounts of fucosylated structures and structures with higher branching (Fig. 5A–E). Moreover, the secreted N-glycomes of both EMT and non-EMT cells comprised of primarily complex-type N-glycans (>80%). We also found one paucimannose structure (peak 3). However, relative ratios of specific glycan structures were dramatically different resulting in N-glycome profiles specific for each cell line (Fig. S5). This difference was expected because of different expression profiles between EMT and non-EMT cells and thus the different composition of secretomes. Besides, it was demonstrated that breast cell lines exhibit specific N-glycosylation signatures of their secretomes relative to their tumorigenic level and breast cancer subtype (Lee et al. 2014).

**Figure 5.**
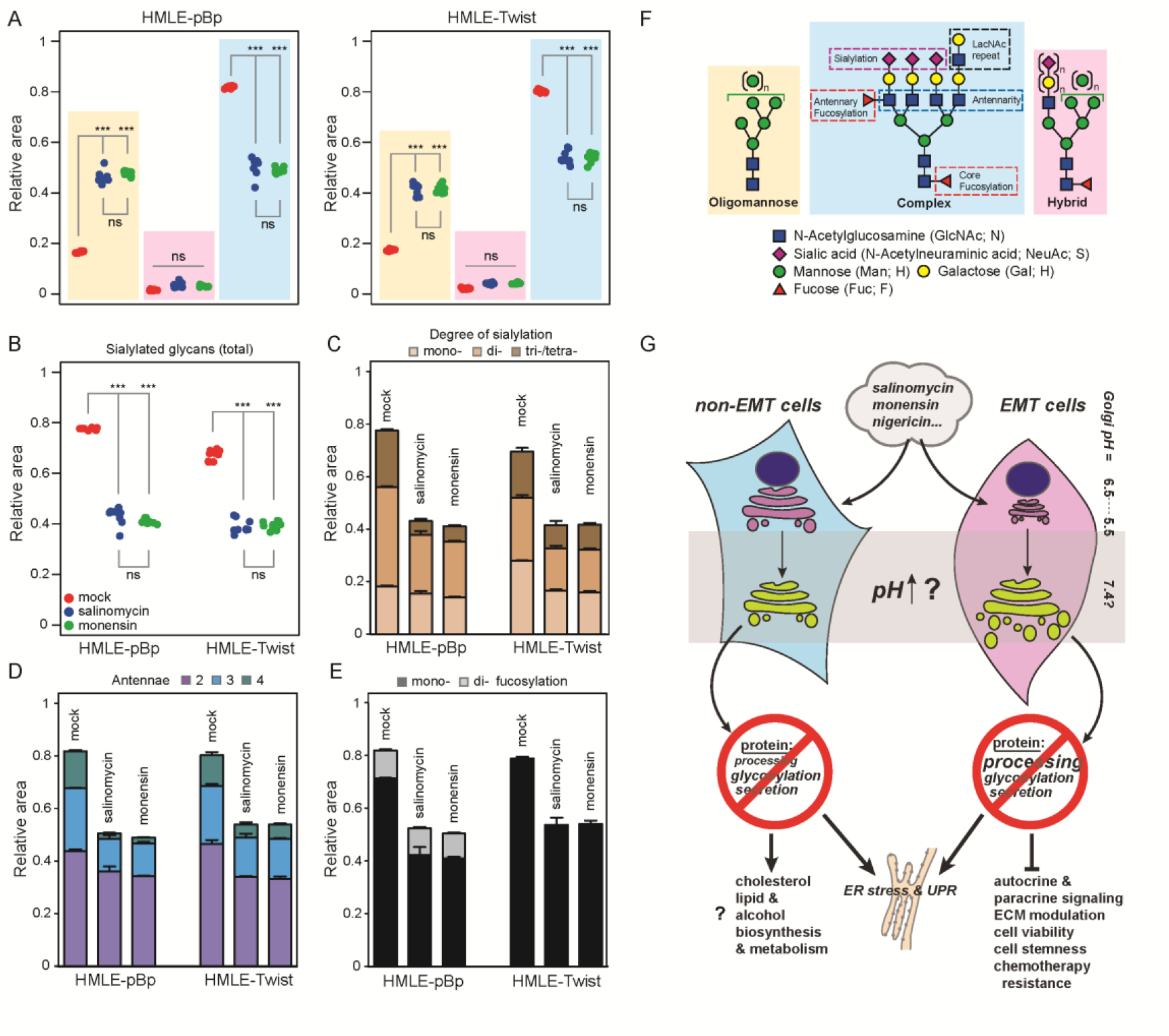
Salinomycin and monensin suppress synthesis of complex N-glycan structures of secreted proteins. (A) Comparative analysis of N-glycans from secreted proteins of mock, Sal and Mon treated non-EMT (HMLE-pBp, left panel) and EMT cells (HMLE-Twist, right panel). The glycans were separated into oligomannose, hybrid and complex glycans (shading box colors follow the scheme in (F)). Each area under the peak from chromatograms (Fig. S6) was integrated and its relative amount for each glycan type was summed and plotted (n = 9, 3 technical replicates from each of 3 biological experiments). Total relative amount of sialylated glycans (B) and degree of sialylation by number of sialic acid residues present on each glycan structure (C). (D) Distribution of branched glycan structures organized by number of antennas. (E) Degree of fucosylated glycan structures organized by the number of fucose residues present. (F) Scheme of N-glycan structures organized by glycan types: oligomannose, hybrid and complex. Background color corresponds to glycan type in A and Figure S6D. (G) Proposed mechanism of ionophores’ selectivity against EMT cells. Ionophores induce exchange of cations for proton, leading to dissipation of H^+^ gradient over the Golgi membrane, which in turn leads to disruption of normal protein processing, glycosylation and secretion. Improperly processed and/or glycosylated proteins accumulate in the ER and the GA inducing ER stress and UPR. On the other hand, abnormal protein trafficking and glycosylation hinders normal cell signaling and communication leading to diminished EMT maintenance, and in non-EMT cells leads to induction of cholesterol, lipid and alcohol biosynthesis. One-way ANOVA with post-hoc Tukey’s multiple comparison test was used for statistical analysis from n = 3 individual experiments (ns – non significant, ***p < 0.001)

After chromatogram integration, we defined 33 distinct chromatographic peaks in EMT and 30 peaks in non-EMT cells (Fig. S5 and Tables S2 and S3). The N-glycan structures present in each peak were determined by HILIC-UPLC-FLR-MS/MS approach. Many peaks contained more than one glycan structure, but in most cases, one single structure was the most abundant (i.e. the major structure; Tables S2 and S3, and Supplementary file S3). Glycan profile of EMT cells was dominated by 3 peaks (14, 16 and 20) that represented 39.3% of the total chromatogram. On the other hand, the glycan profile of non-EMT cells was dominated by 4 peaks (16, 17, 19 and 20) representing similarly 39.2% of the total glycome profile. Major glycans found in peaks 16 and 20 from HMLE-Twist cells matched to major glycans found in peaks 16, 17 and 19, 20 from HMLE-pBp cells (FA2G2S1 and FA2G2S2, respectively; Fig. S5D and Tables S2 and S3). Further, EMT cells lacked terminal fucosylation reflecting the fucosyl transferases’ expression (Supplementary file S2A). In addition, EMT cells had reduced total relative amount of sialylated glycans in comparison to non-EMT cells. Besides, sulfated or phosphorylated N-glycans (S/P) were found only in EMT cells, which had 4 such structures forming individual peaks 17, 21, 28 and 32 (Fig. 5A–E, S5, and Tables S2 and S3).

Treatment with either Sal or Mon had a drastic impact on N-glycosylation of secreted proteins in both cell lines. Strikingly, the effect of these two ionophores was extremely similar. Detailed analysis showed that each observed change was in the same direction and of similar potency when compared to mock-treated samples (Fig. 5A–E, S5B–D, and Tables S2 and S3). Thus, as treatments with both ionophores produced N-glycome profiles that were hardly distinguishable, we will mainly discuss treatment vs. control regarding glycosylation. The differential effect of the treatment between cell lines was marginal (Fig. 5A and S5D). Treatment did not completely abolish N-glycosylation of secreted proteins. Rather, it mostly blocked the processing of larger and more complex N-glycan structures as the relative amount of these glycans was significantly reduced in both cell lines. In turn, this led to a relative increase of simple, oligomannose-type N-glycans structures, reflecting the semi-stalled glycosylation machinery (Fig. 5A and S5D). Moreover, an increase in oligomannose structures after treatment is indicative of GA rather than ER -specific action, because mannoses are formed at ER. Therefore, a decrease in oligomannose structures would be expected if the treatment is ER - specific. A similar phenotype was observed on all structures that appear with increased N-glycan complexity. For instance, we observed decreased total sialylation, with especially affected tri-/tetra-antennary sialylated structures; decreased branching and decreased fucosylation of the N-glycome (Fig. 5B–E). These findings demonstrated that Sal and Mon disturb N-glycosylation at the stage of building complex glycans, which takes place within the GA.

## Discussion

Salinomycin has been in the spotlight of cancer research ever since its EMT/CSC selective capabilities were revealed (Gupta et al. 2009). While numerous effects of Sal in different cellular models were reported, the mechanism of its selectivity against EMT cells remained elusive. Our work, for the first time, establishes Sal as a Golgi apparatus disturbing agent, positioning it along with established GA inhibitors, monensin and nigericin. The EMT/CSC selectivity of Mon was only recently experimentally confirmed in a prostate cancer model (Vanneste et al. 2019). In our experiments, both Sal and Mon lead to marked morphological perturbations of the GA accompanied by hindered post-translational protein modifications, elevated ER-stress and UPR, diminished secretion and highly altered N-glycosylation of secreted proteins. These changes induce the expression of the ER, the GA, and membrane-related genes, pointing to problems in the secretory pathway. We hypothesize that Sal leads to the alkalization of the GA via increased leakage of protons from the organelle in exchange for K^+^ ions. Increased pH then leads to changes similar to the effect of monensin and nigericin (Fig. 5G) (46,47) since the maintenance of acidic compartments of the secretory pathway is crucial for proper protein processing, glycosylation, and sorting, as well as the organelle morphology (Weisz 2003; Rivinoja et al. 2012). Moreover, the activation of cadherins requires proteolytic removal of their prodomain in the late GA compartments (TGN) by proprotein convertases, most probably Furin (Koch et al. 2004; Maret et al. 2010). We observed incomplete processing of N-cadherin in EMT cells after Sal and Mon. Likewise, Mon was shown to exert its best ionophoric activity in the cholesterol-rich membranes (trans-Golgi) as compared to the low-cholesterol membranes (cis-Golgi) (Orci et al. 1981; Dinter and Berger 1998). In addition, we primarily detected diminished relative amounts of complex N-glycan structures after the treatment with Sal and Mon. All glycosyltransferases that introduce and process components of complex N-glycans (e.g. MGATs – branching, STGALs – sialylation and FUTs – fucosylation) are found within cisternae of the GA, mostly in TGN (Varki et al. 2009). Taken altogether, our results clearly demonstrate that Sal specifically targets the GA and position it as a GA disturbing agent.

Sal targeted the GA in both EMT and non-EMT cells. However, particular downstream effects resulted in differential outcomes in EMT and non-EMT cells. For example, Sal induced the expression of the ER- and the GA-related genes in both cell lines, but at the same time the activation of biological processes in EMT and non-EMT cells differed substantially. Likewise, Sal induced severe fragmentation of the Golgi cisternae only in EMT cells. Also, the removal of E-cadherin prodomain in non-EMT cells (HMLE-pBp, HMLE-shGFP or MCF-7), which requires same proprotein convertases, was not inhibited by Sal. Thus, Sal and Mon diversely affect the GA in regard to their EMT status, and/or at different points within this organelle. This, in turn, selectively affected viability of EMT cells, most probably by abrogating their proper maintenance. For example, changes in glycosylation are of major importance for understanding the significance of the EMT during carcinogenesis (Taniguchi and Kizuka 2015). Interestingly, the effect on N-glycosylation in EMT and non-EMT cells was very similar, contrary to differences detected in gene expression, cadherin maturation, and the GA morphology. This suggests a higher dependence of EMT cells on the processes executed in the GA, positioning it as their sensitive spot. However, we measured only the effect on N-glycosylation of secreted proteins. It was demonstrated that the requirements for glycosylation appear to be more stringent for membrane than for secreted proteins (Fiedler and Simons 1995). Of note, we can assume that aberrations of glycan structures extend to non-secreted and membrane proteins, and also to additional glycosylation classes processed within the GA (e.g. O-linked and lipid glycosylation, glucosaminoglycans), which might be the underlying reason for different sensitivity of EMT vs. non EMT-cells.

Furthermore, we observed the activation of ER-related UPR, which was already established as the mechanism behind the selectivity towards EMT cells (Feng et al. 2014). The activation of UPR in our experimental setting is likely a downstream effect of inducing the GA-stress through the excessive accumulation of properly folded proteins. It was previously shown that the inhibition of N-glycosylation inside the GA leads to the accumulation of improperly glycosylated proteins within the GA, the diminished secretion of proteins and the induction of the ER-stress related UPR (Xu et al. 2010). In our study, Sal activated PERK and not the IRE1 branch of UPR, positioning it in different cluster of UPR activating compounds together with Mon and brefeldin A, contrary to the cluster containing tunicamycin and thapsigargin (Shinjo et al. 2013). These observations led us to conclude that EMT cells are sensitive to Golgi disturbing agents and to propose the GA as a valid target against cancer cells with EMT properties.

Given the close physical and functional connection between the GA and the ER, it is surprising that the Golgi complex has remained a largely unexplored drug intervention target. The finding that EMT cells are sensitive to Golgi disturbing agents has an important implication for the treatment of highly malignant tumors, especially the ones with high proportions of EMT cells. The ionophores used in this study target multiple aspects of the GA function and are known to carry diverse toxic effects to the whole organism (Boehmerle et al. 2014). Albeit, some have shown promising pre-clinical efficacy against therapy-resistant cancers (Dewangan et al. 2017). Further studies focused on elucidating the mechanisms that regulate the GA function may lead to therapeutic targeting of its discrete components. For example, narrowing down the specific GA-related process could propose more precise inhibitors (e.g. against a specific glycosyltransferase) with better efficacy and lower toxicity.

## Methods

### Cell culture and reagents

HMLE cells expressing Twist or empty vector (HMLE-Twist and HMLE-pBp), as well as expressing shRNA targeting E-cadherin or GFP (HMLE-shEcad and HMLE-shGFP) and SUM 159 were a kind gift from Dr. Robert A. Weinberg’s lab. HMLE cells were maintained in a 1:1 mixture of HuMEC Ready Medium (Gibco) and DMEM complemented with 10% FBS, 2 mM L-glutamine, 100 U/mL penicillin, 100 μg/mL streptomycin, 2.5 μg/ml insulin and 0.5 μg/mL hydrocortisone (Sigma-Aldrich). MCF-7 and MDA-MB-231 cell lines were obtained from ATCC and maintained in DMEM complemented with 10% FBS, 2 mM L-glutamine, 100 U/mL penicillin and 100 μg/mL streptomycin, while SUM 159 cells were grown in Ham’s F12 media with 5% FBS, 1 μg/ml hydrocortisone and 5 μg/ml insulin (Sigma-Aldrich). All cells were grown in a humidified atmosphere at 37°C with 5% CO2. EMT and non-EMT status was regularly checked by probing for EMT markers (CD24/CD44 by FACS and E- and N-cadherin by western blot). Salinomycin (S4526), nigericin (N7143), monensin (46468), paclitaxel (T7402) and tunicamycin (T7765), brefeldin A (B6542) and GalNAc(α)-O-bn (B4894) were purchased from Sigma-Aldrich.

### Proliferation assays

#### MTT

Cells were seeded (3000 per well) in standard 96-well microtiter plates and left to attach. Next day, test compounds were added at 10 different concentrations in quadruplicates. The cell growth rate was evaluated after 72 hours of incubation by adding MTT reagent (Sigma-Aldrich). Obtained results for each compound were calculated and normalized to mock-treated cells. Dose-response curves were plotted in GraphPad Prism using non-linear fit.

#### Flow Cytometry

For mixed culture experiments, cells were seeded in 6-well culture plates at 1:1 ratio (75000 EMT and 75000 non-EMT cells) and left to attach. Next day, media was replaced by media containing vehicle (DMSO) or tested compound. After 72 hours of treatment, cells were detached, washed in blocking solution (2% BSA in PBS) and immunolabeled with anti-CD24 PerCP/Cy5.5 and anti-CD44 FITC primary antibodies (BioLegend, 1:50, 311115 and 1:100, 338803, respectively) for 30 minutes at RT. Cells were then washed twice with PBS and transferred to FACS tubes. 10000 cells were analyzed on a BD FACSCalibur, and percentages of CD24/CD44 specific populations were calculated in FlowJo (TreeStar, Inc.).

### Western blot

Cells were collected by trypsinization, pelleted and washed with PBS on ice. Pellets were lysed with RIPA buffer supplemented with complete protease inhibitor (Roche) supplemented with phosphatase inhibitors (NaF and Na3VO4, Sigma-Aldrich) and subsequently sonicated using Bioruptor (Diagenode). Total concentration of proteins in cell lysates was measured by Pierce BCA Protein Assay Kit (Thermo Fisher Scientific) according to the recommendations from the manufacturer. 50 μg of protein per sample were loaded on a gel, separated using SDS-PAGE and transferred to PVDF membrane (Carl Roth). After protein transfer, membranes were stained using naphthol blue solution (0.1% naphthol blue in 10% methanol, 2% acetic acid) to visualize transferred proteins that was used to evaluate equal sample loading. Next, membranes were incubated in blocking solution (5% non-fat milk in TBST) for 20 minutes at room temperature. After blocking, membranes were incubated with primary antibodies at 4°C overnight. Primary antibodies used: E-cadherin (1:500, Becton-Dickinson, 610181), N-cadherin (1:500, Becton-Dickinson, 610920), proN-cadherin (1:500, R&D Systems, AF1388), EGFR (1:1000, Cell Signaling Technology, 4267), TFRC (1:500, Thermo Fisher Scientific, 13-6800), eIF2α (1:1000, Cell Signaling Technology, 9722), p-eIF2α (1:1000, Cell Signaling Technologies, 3597), CREB-2 (1:100, SCBT, sc-200). After washing the membranes with TBST, membranes were incubated with complementary HRP conjugated secondary antibody (sheep anti-mouse IgG-HRP, 1:10000, GE Healthcare, NA931; goat anti-rabbit IgG-HRP, 1:5000, Bio-Rad, 1706515; rabbit anti-sheep IgG-HRP, 1:3000, Invitrogen, 61-8620) for 2 hours at room temperature and subsequently washed again with TBST before detection of signal by Western Lightning Plus-ECL reagent (Perkin-Elmer, USA). Emitted signal from membranes was collected and visualized with UVITEC imaging system (Cleaver Scientific Ltd) and final images were prepared in Photoshop CS2 and Illustrator CS2 (Adobe). To remove N-glycans from proteins, cell lysates were additionally treated with 16 units of PNGase F (New England Biolabs) per μL of lysate complemented with 1% NP-40. De-glycosylation reaction was performed at 37°C for 60 minutes.

### Microscopy

Cells were seeded onto poly-L-Lysine treated glass cover slides, left to attach and the next day treated with compounds as indicated. Cellular compartments were stained according to the following protocol: After 24 hours of incubation slides were fixated with 4% PFA in PBS for 10 minutes and then permeabilised with blocking solution (2% BSA, 0.05% Tween-20 in PBS) for another 10 minutes. Slides were then incubated with primary antibody in blocking solution for 30 minutes at room temperature (RT). Primary antibodies used: GM130 (1:500, BD Biosciences, 610822), GIANTIN (1:2000, Abcam, ab24586), ERGIC53 (1:1000, Enzo Life Sciences, ALX-804-602) and β’COP (1:500, rabbit polyclonal anti-β’COP antibody was a kind gift from Dr. Rainer Pepperkok, EMBL Heidelberg). Slides were then washed and incubated with complementary goat anti-rabbit Alexa Fluor 488, goat anti-mouse Alexa Fluor 568 or goat anti-mouse Alexa Fluor 488 secondary antibody (Thermo Fisher Scientific, A-11034, A-11031 and A-11001, respectively) diluted 1:300 in blocking solution at RT. In the end, slides were extensively washed with blocking solution, and in the last washing step 100 ng/mL of DAPI was used as a counter stain for DNA. Slides were then mounted onto glass slides using Fluoromount antifade reagent (Sigma-Aldrich). Images were collected on a Leica SP8 confocal microscope, processed in ImageJ and prepared for publication in Illustrator CS2 (Adobe).

### RNA-Seq

Cells were seeded onto 10 cm plates (1.3 million cells per plate) and left to attach overnight. Next day, media was replaced by media containing vehicle (DMSO) or 0.2 μM Sal. Ater 24 hours, total RNA from biological triplicates of HMLE-Twist and HMLE-pBp cells was extracted (12 biological samples in total) with Quick-RNA Miniprep kit (Zymo Research) according to manufacturer protocol. DNase I was used to remove possible DNA contamination. RNA concentration was measured on Qubit 3.0 Fluorometer (Thermo Fisher Scientific) and the quality of RNA was additionally checked with Bioanalyzer 2100 (lowest RIN = 9.7, Agilent). cDNA library was prepared with TrueSeq Stranded mRNA kit (Illumina) from 750 ng of RNA (average cDNA size was ∼345bp). Library was sequenced on NextSeq500 (Illumina) using high output flow-cell in paired-end mode (75 + 75 cycles). Raw data yielded over 690 million high-quality reads (Q30 > 75%). Reads were uploaded to BaseSpace Sequence Hub (Illumina) and aligned to human reference genome (hg19) via RNA-Seq Alingment app (STAR alignment, ver 1.1.1). Differential expression of genes after treatment for each cell line and between EMT and non-EMT cell lines was executed via DESeq2 app (ver 1.1.0) in paired mode. The data discussed in this publication have been deposited in NCBI’s Gene Expression Omnibus (Edgar et al. 2002) and are accessible through GEO Series accession number (GSE124975).

### Gene Ontology and GSEA

#### GO

For Gene Ontology analysis we used Gene Ontology Enrichment Analysis and Visualization tool (GOrilla, http://cbl-gorilla.cs.technion.ac.il/) by loading single ranked genes from each differential gene expression experiment (genes were ranked according to differential expression from highest to lowest). Additionally, we performed GO analysis of overlapping genes from different experimental settings with Panther GO algorithm (http://www.geneontology.org/). Venn diagrams were plotted via web tool of Ghent University (http://bioinformatics.psb.ugent.be/webtools/Venn/).

#### GSEA

For gene set enrichment analysis (GSEA), expression data obtained after treatment with Sal in both EMT and non-EMT cells was analyzed for enrichment of top 100 genes induced by monensin (GSE5258), nigericin and HyT36 Golgi UPR (Serebrenik et al. 2018) (GSE99490), doxorubicin (GSE39042), 4MU-xyloside (GSE117938), brefeldin A (GSE94861), tunicamycin and thapsigargin (GSE24500). In addition, GSEA was performed for genes in gene ontology category Glycosylation downloaded from Molecular signature database (MsigDB). All GSEA analyses were executed with the use of GSEA software downloaded from Broad Institute (MIT)(Subramanian et al. 2005).

### Elucidation of additional EMT selective compounds

Raw data from initial compound screen by Gupta et al. (6) was downloaded and analyzed in R. To generate additional compounds with selective properties we performed 3 algorithms, which yielded 32, 54 and 183 compounds combining to total of 227 unique compounds. Algorithms: 1) shGFP BsubvalueA (or B) > of mean of BsubvalueA (or B) AND shEcad ZscoreA (or B) is < mean – 1 s.d. of ZscoreA (or B); 2) shGFP BsubvalueA (or B) > of mean of average (BsubvalueA, BsubvalueB) AND shEcad ZscoreA (or B) is < mean – 1 s.d. of ZscoreA (or B); 3) shGFP BsubvalueA (or B) > of mean-1 s.d. of BsubvalueA (or B) AND shEcad ZscoreA (or B) is < mean – 2 s.d. of ZscoreA (or B).

### RT-PCR

For the analysis of UPR-related splicing of XBP1, 150000 cells were seeded in 6-well plates and left to attach. Next day, media was replaced with media containing vehicle (DMSO) or tested compounds. After 24 hours of treatment, cells were washed with ice-cold PBS and RNA was isolated from cells using TRIzol reagent (Sigma). Reverse transcription and subsequent PCR amplification was done with OneTaq RT-PCR Kit (New England Biolabs). Primers: GAPDH-For (gagtcaacggatttggtcgt), GAPDH-Rev (ttgattttggagggatctcg), XBP1-For (aaacagagtagcagctcagactgc) and XBP1-Rev (tccttctgggtagacctctgggag).

### ELISA

Cells were grown in 24-well plates (75000 cells/well) in 1 mL media containing vehicle (DMSO) or compounds. After 24 hours, media above cells was collected and analyzed for the amount of IL-8 secreted. ELISA was performed with Human IL-8 ELISA Ready-SET-Go kit (Affymetrix eBioscience) according to manufacturer’s protocol.

### Nile Red staining

Cells were seeded onto poly-L-Lysine treated glass cover slides, left to attach and the next day treated with compounds as indicated. Lipid droplets were stained according to the following protocol: After 24 hours of incubation slides were washed with PBS twice and then fixated with 4% PFA in PBS for 15 minutes at RT. After fixation, cells were again washed twice with PBS and then stained with 150 nM Nile Red (Sigma-Aldrich, 72485) in PBS for 15 minutes at RT. Cells were then washed with PBS twice and mounted onto microscopy slides using Fluoromount antifade reagent containing 100 ng/mL DAPI (Sigma-Aldrich). Images were collected on a Leica SP8 confocal microscope, processed in ImageJ and prepared for publication in Illustrator CS2 (Adobe).

### Measuring cytosolic calcium

#### Flow cytometry

500000 cells in suspension were loaded with 3 μM Fluo-4 AM (Thermo Fisher Scientific, F14201) in DMEM without serum supplemented with 0.02% Pluronic-F127 (Sigma, P2443) for 60 min at 37°C. The cells were then washed in HBSS and left in DMEM without serum for additional 20 minutes at 37°C to allow complete de-esterification of intracellular Fluo-4 AM. Cells were then washed twice with HBSS and, after the last wash a 400 μL of compound in HBSS was added to cells (final concentration as indicated). Green fluorescence of 10000 cells was measured 10 and 30 minutes after the addition of compound on a BD FACSCalibur. The collected data was further analyzed in FlowJo (TreeStar, inc.).

#### Confocal imaging

25000 cells were seeded on a 3.5 cm dishes with glass bottom (IBIDI, 81156) and left to attach overnight. On the day of imaging, cells were loaded with Fluo-4 AM as for flow cytometry above. For fluorescence imaging, cells were washed twice with HBSS and 1.2 mL was left over cells for imaging. After desired area with cells was found, cells were imaged every 15 seconds for 10 minutes. 400 μL of 4× concentration of compound in HBSS was added directly to the cells (final concentration indicated) under the microscope before the second image (after ∼10 seconds). Images were collected on a Leica SP8 confocal microscope with 40× immersion objective. The Fluo-4 fluorescence intensity of multiple cells was measured in ImageJ and normalized to the same cell from the first image.

### Analysis of mitochondrial depolarization

500000 cells in DMEM were treated with salinomycin at indicated concentrations. After 30 minutes of incubation 2.5 μg/mL of JC-1 mitochondrial potential fluorescent probe (eBioscience, EBI-65-0851-38) was added to the cells for additional 30 minutes of incubation (1h total of Sal treatment). Cells were then placed on ice and analyzed on a BD FACSCalibur flow cytometer, and percentages of red and green fluorescent specific populations were calculated in FlowJo (TreeStar, Inc.).

### Analysis of the N-Glycome on secreted proteins

Cells were seeded onto 10 cm plates (1 million cells per plate) and left to attach overnight. Next day, media was discarded, cells were washed with PBS 4 times and 9 mL of media (without FBS or supplements) containing vehicle (DMSO) or compound was added to each plate. After 48 hours, plates were placed on ice, media was collected into tubes and residual cells and large debris was pelleted at 2000g. Supernatant was collected and concentrated with Amicon Ultra centrifugal filter devices with 10 kDa cut-off membrane (Merck). Protein concentration was measured by Pierce BCA Protein Assay Kit (Thermo Fisher Scientific) according to the recommendations from the manufacturer. 15 μg of proteins from each sample were denatured with the addition of SDS and by incubation at 65 °C. The excess of SDS was neutralized with Igepal-CA630 (Sigma-Aldrich) and N-glycans were released following the addition of PNGase F (Promega) in PBS. The released N-glycans were labeled with 4-amino-N-(2-diethylaminoethyl) benzamide (Procainamide; ProA). Free label and reducing agent were removed from the samples using hydrophilic interaction liquid chromatography solid-phase extraction (HILIC-SPE). Fluorescently labeled N-glycans were analyzed by HILIC-UPLC-FLR-MS/MS on an Acquity UPLC instrument (Waters) consisting of a quaternary solvent manager, sample manager, and an FLR fluorescence detector set with excitation and emission wavelengths of 310 and 370 nm, respectively. N-glycans were separated on a Waters BEH Glycan chromatography column, 150 × 2.1 mm i.d., 1.7 μm BEH particles, with 100 mM ammonium formate, pH 4.4, as solvent A and ACN as solvent B. The separation method used a gradient of 32-38% solvent A for 16 min and 38%-55% solvent A for next 24 min. Flow rate of 0.56 mL/min was constant during the whole analytical run. Samples were maintained at 10 °C before injection, and the separation temperature was 25 °C. The UPLC was hyphenated to the Bruker compact Q-Tof mass spectrometer (MS), both controlled using HyStar software (Bruker) version 4.1. UPLC was coupled to MS via Ion Booster (Bruker) ion source. Capillary voltage was set to 2250 V with nebulizing gas at pressure of 5.5 Bar. Drying gas was applied to source at a flow rate of 4 L/min and temperature of 300 °C, while vaporizer temperature was set to 300 °C and flow rate of 5 L/min. Nitrogen was used as a source gas, while argon was used as collision gas. Ion energy was set to 5 eV, transfer time was 100 μs. Spectra were recorded in mass range from 50 m/z to 4000 m/z at a rate of 0.5 Hz. For MS acquisition collision energy was set to 4 eV. Fragment spectra were acquired using auto MS/MS mode. HILIC-UPLC-FLR chromatograms were used for quantification, and abundance of each glycan was expressed as percentage of total integrated area. N-glycan compositions and structural features were assigned using GlycoMod (ExPASy) (Cooper et al. 2001) (http://web.expasy.org/glycomod/) and GlycoWorkbench (Ceroni et al. 2008).

### Statistics

All graphics with error bars are presented as mean ± s.d. except Fig. S3D where mean ± s.e.m. was plotted. To determine statistical significance between samples, one-way ANOVA with Dunnett’s (Fig. 3C and D, Fig. S3B) or Tukey’s (Fig. 5 and Tables S2 and S3) multiple comparison post-hoc test was used. Statistical calculations were performed in GraphPad Prism and generation of graphics in Excel, R Studio, and Adobe Illustrator CS2 (NS – non-significant; * - p < 0.05, ** - p < 0.01 and *** - p < 0.001).

## Acknowledgments

The authors thank Dr. Robert A. Weinberg and Dr. Tamer T. Onder for providing us with HMLE and SUM 159 cell lines. We thank Dr. Rainer Pepperkok and his lab for help, expertise and shared reagents regarding microscopy of cellular compartments. We are also thankful to Dr. Siniša Volarević and his lab for UPR reagents, Dr. Oliver Vugrek and Filip Rokić for help with RNA-Seq experiment, Lucija Horvat for assistance with confocal microscopy, and Dr. Fran Supek for critical reading of the manuscript.

## Statements and Declarations

### Author Contributions

Marko Marjanović, Ana-Matea Mikecin Dražić and Marijeta Kralj contributed to the study conception and design. Experimental work, analysis of the data, visualization of the data, review and editing of the manuscript were performed by Marko Marjanović, Ana-Matea Mikecin Dražić, Marija Mioč, Filip Kliček, Mislav Novokmet. The first draft of the manuscript was written by Marko Marjanović and all authors commented on previous versions of the manuscript. All authors read and approved the final manuscript.

### Data availability

The datasets generated during the current study are available in the NCBI’s Gene Expression Omnibus and are accessible through GEO Series accession number (GSE124975), or are included in this published article and its supplementary information files.

