## Supporting information for "Salinomycin disturbs Golgi apparatus function and specifically affects cells in epithelial-to-mesenchymal transition"

[a] Division of Molecular Medicine, Ruđer Bošković Institute, Bijenička c. 54, 10000 Zagreb, Croatia
[b] GENOS, Glycoscience Research Laboratory, Borongajska c. 83, 10000 Zagreb, Croatia
[c] These authors contributed equally
[d] ORCID: 0000-0001-9850-0561

[e] ORCID: 0000-0001-5290-6945

[f] ORCID: 0000-0002-7165-2722

**This PDF file includes:**

Figures S1 to S5

Tables S1 to S3

Legends for Files S1 to S3

**Other supplementary materials for this manuscript include the following:**

Files S1 to S3 as separate excel files.

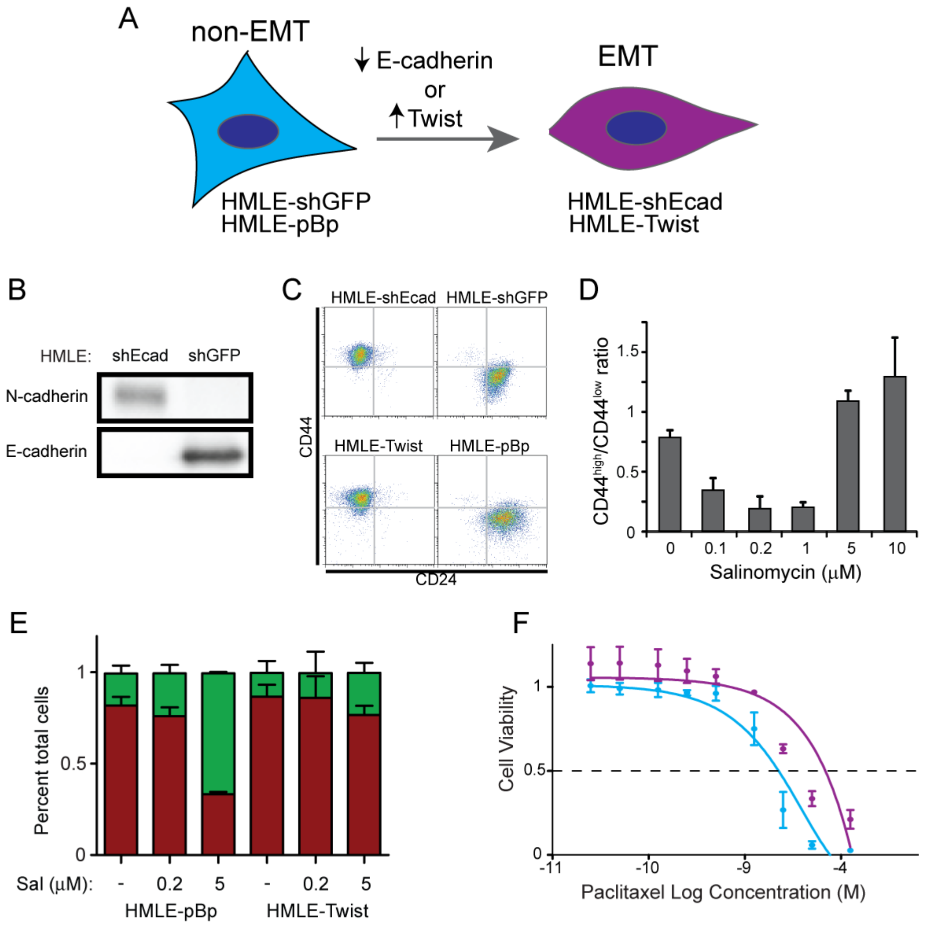


**Figure S1. Golgi disturbing agents are selective towards EMT cells.**

(A) EMT model used in the study.

(B) Western blot analysis of N- and E-cadherin protein expression (EMT and non-EMT marker, respectively) in HMLE-shEcad and HMLE-shGFP cells.

(C) Representative CD24/CD44 flow cytometry profiles of HMLE-shEcad and HMLE-shGFP, HMLE-Twist and HMLE-pBp (EMT and non-EMT cell pairs).

(D) HMLE-shEcad and HMLE-shGFP were seeded on day 0 at 1:1 ratio and then treated with salinomycin (Sal) at increasing concentrations. Ratio of $CD44^{high}/CD44^{low}$ (EMT/non-EMT) cells after 3 days of treatment of mixed culture is shown.

(E) High concentration of Sal depolarizes mitochondria of non-EMT cells. Cells were treated with Sal for 1h and then stained with JC-1. Red (polarized) and green (depolarized) fluorescence was measured by flow cytometery and their respective percentages were calculated.

(F) Dose-response curves of HMLE-shEcad (purple, EMT) and HMLE-shGFP (blue, non-EMT) breast cells treated with paclitaxel. MTT assay was performed after 72 hours of treatment.

Data are presented as mean ± s.d., n = 3 from duplicates in D and E, n = 3 from quadruplicates in F.

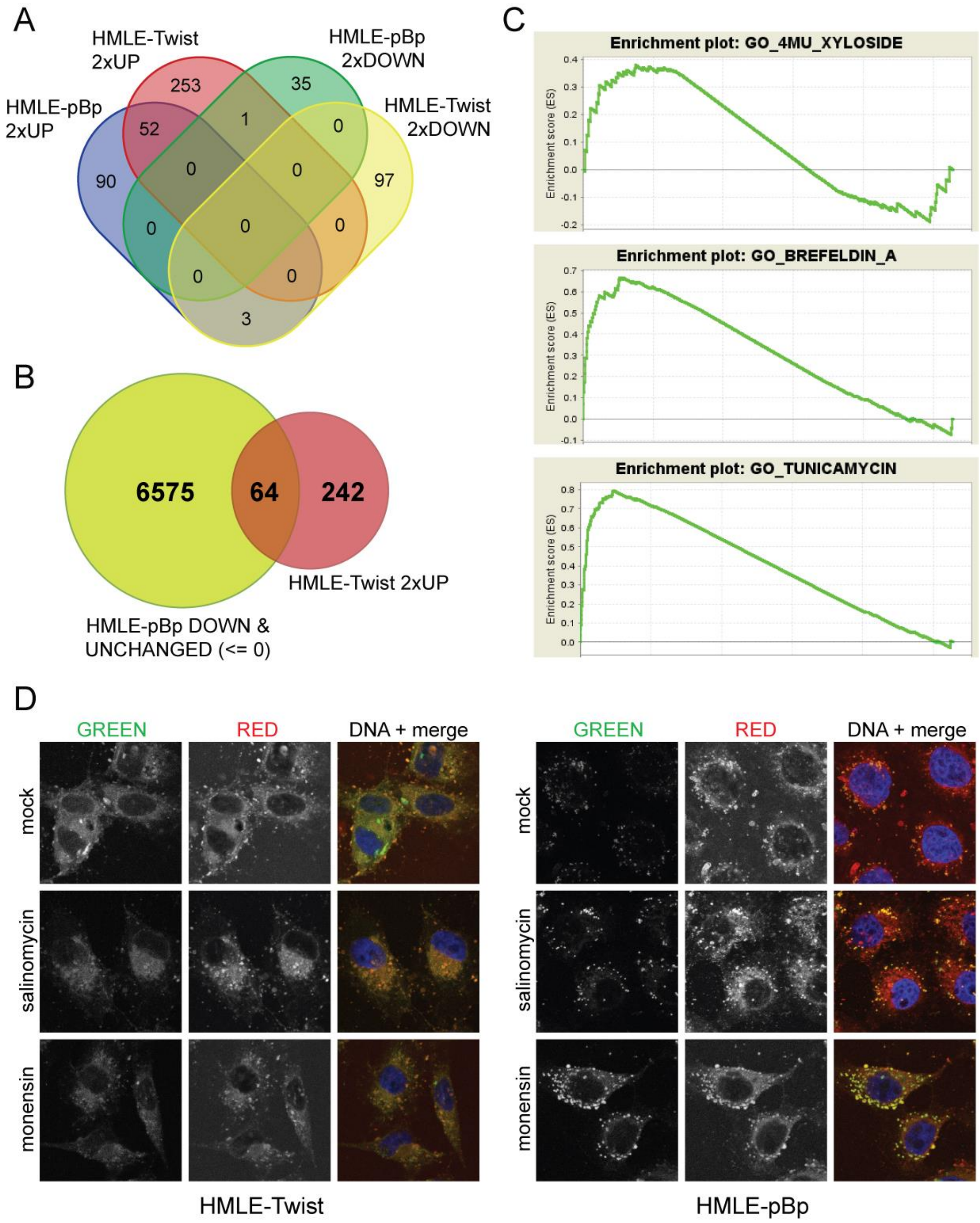


**Figure S2. Differential gene expression, gene ontology and GSEA analysis of RNA-seq experiment.**

(A and B) HMLE-Twist and HMLE–pBp were treated with 0.2 µM salinomycin for 24 hours and RNA-seq experiment was performed. Venn diagram representation of number of differentially expressed genes is presented. Direction and threshold of fold change used for the analysis is indicated.

(C) GSEA analysis for gene sets of known Golgi (brefeldin A and 4MU-xyloside) and ER stressor (tunicamycin) on RNA-seq data from HMLE-Twist cells treated with salinomycin is presented (analysis was performed on data ordered from largest to smallest, see Methods for details about gene sets used).

(D) Salinomycin and monensin lead to accumulation of cholesterol enriched lipid droplets in non-EMT cells. Representative confocal microscopy images of HMLE-Twist and HMLE-pBp cells after treatment with salinomycin or monensin (0.2 µM) for 24 hours. Cells were stained with Nile Red and red (ex. 565 nm, em. > 635 nm) and green/yellow (ex. 488 nm, em 520-550 nm) fluorescence was collected. DNA was counterstained with DAPI (blue). Maximum projection of 4 slices is displayed.

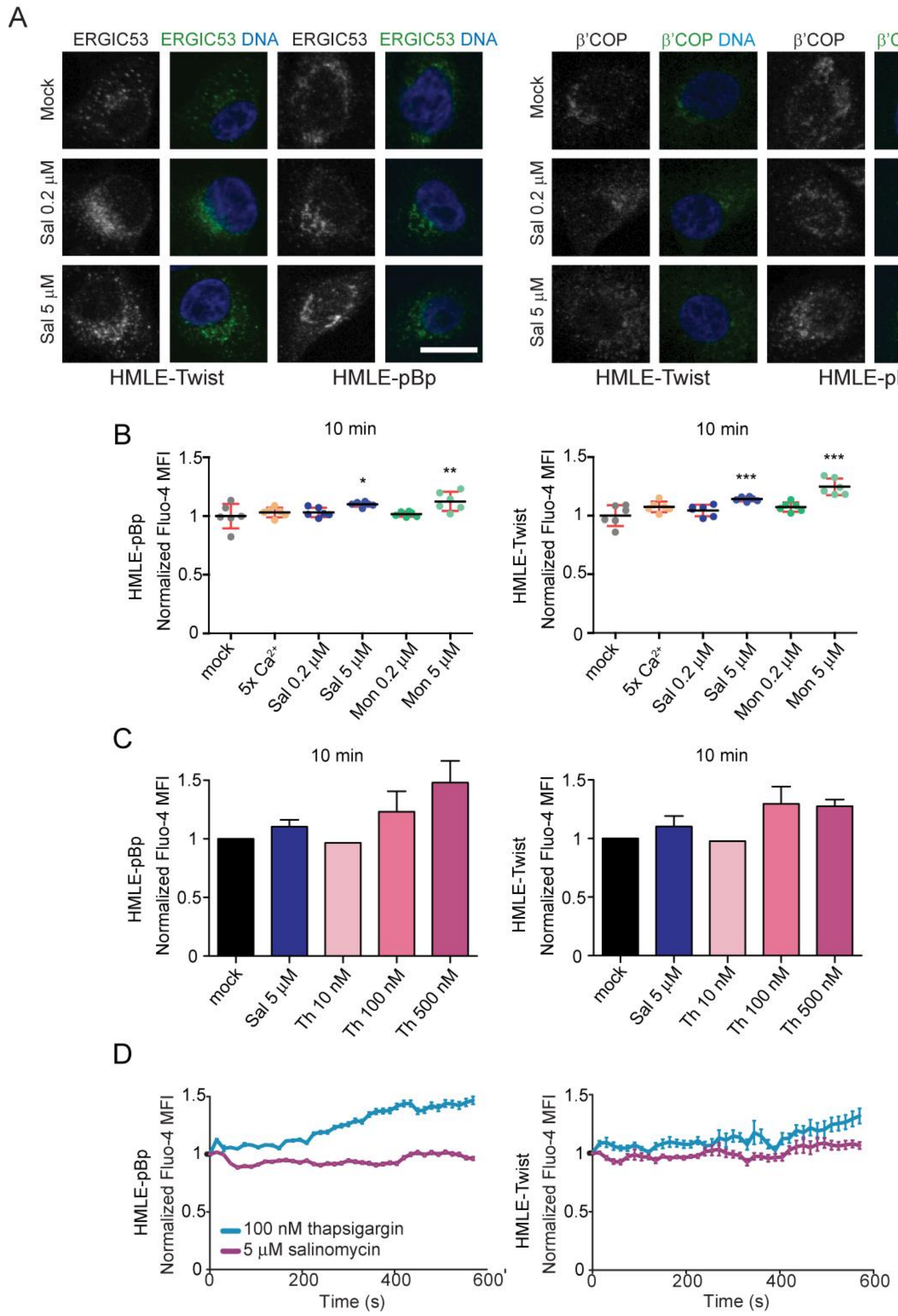


**Figure S3. Salinomycin affects ER-to-Golgi compartment and does not lead to a massive depletion of ER-$Ca^{2+}$ storage.**

(A) Representative confocal microscopy images of HMLE-Twist and HMLE-pBp cells stained with anti-ERGIC and anti-β'COP primary and secondary Alexa Fluor 488 antibodies (green). DNA was counterstained with DAPI (blue). Maximum projection of 10 slices is displayed. Scale bar = 10 μm.

(B and C) Increase in cytosolic calcium levels was monitored as an increase in fluorescence of a $Ca^{2+}$ specific probe (Fluo-4 AM) by flow cytometry after the addition of compounds (Sal – salinomycin, Mon – monensin, Th – thapsigargin). Mean ± s.d. is plotted from n = 6 in (B) and n = 3, 3, 1, 3 and 2 from left to right respectively in (C).

(C) Live imaging of cells loaded with Fluo-4 AM. Confocal microscopy images were taken every 15 seconds, compounds were added on second 10 before acquiring second image. Mean ± s.e.m. is plotted from n = 23 cells (HMLE-pBp) or n = 11 and 9 cells (HMLE-Twist) from one experiment. One-way ANOVA with post-hoc Dunnett's multiple comparison test was used for statistical analysis (*p < 0.05, **p < 0.01, ***p < 0.001)

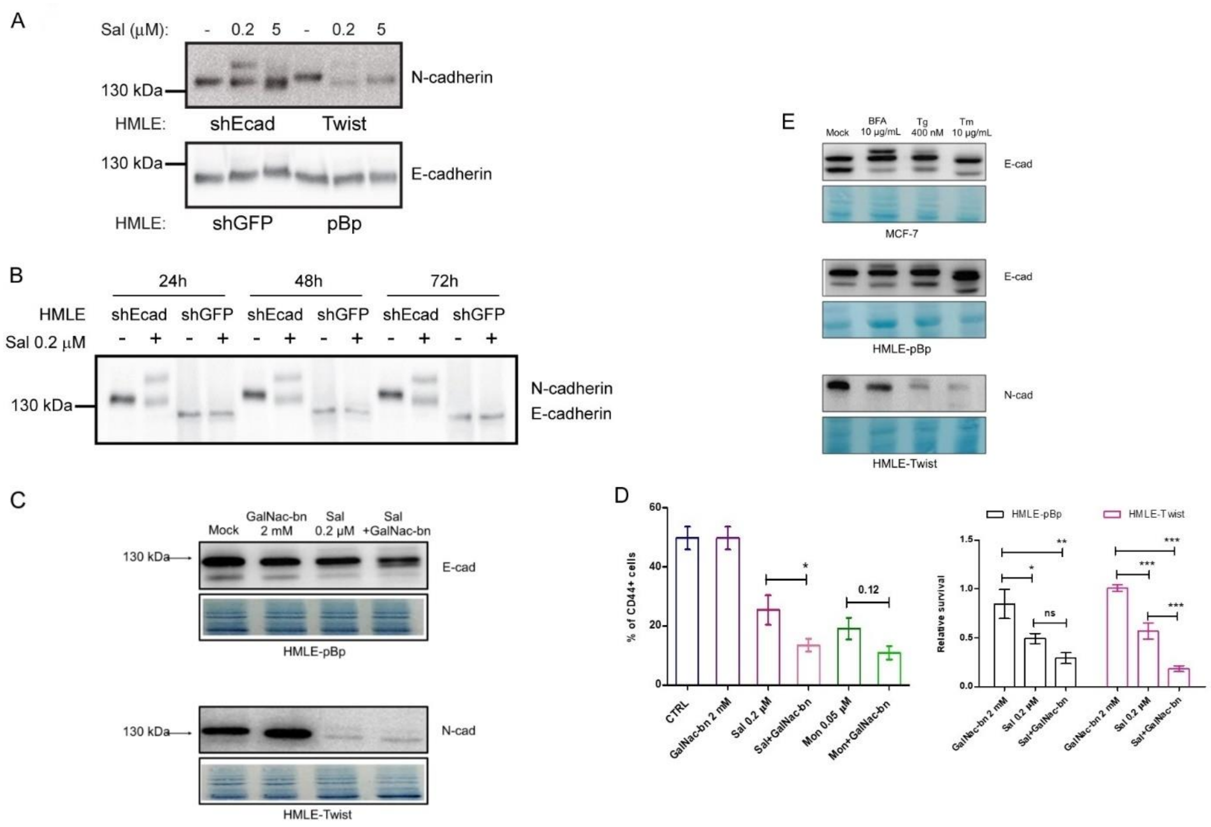


**Figure S4. Salinomycin affects protein post-translational modifications**

(A) The effect of the lower and higher Sal concentrations on N-cadherin processing in HMLE cellular models.

(B) Time-course western blot analysis of N- and E-cadherin after treatment with Sal in HMLE-shEcad and HMLE-shGFP cells.

(C) The effect of GalNAc(α)-O-bn (GalNac-bn) alone or in combination with 0.2 µM Sal on the processing of E- and N-cadherin in HMLE-pBp and HMLE-Twist cells. Amidoblack staining of transferred proteins was used to evaluate equal loading.

(D) Left panel: HMLE-Twist and HMLE-pBp were seeded on day 0 at 1:1 ratio and then treated with salinomycin (Sal), monensin (Mon), GalNAc(α)-O-bn (GalNac-bn) or their combinations at indicated concentrations. Ratio of $CD44^{high}/CD44^{low}$ (EMT/non-EMT) cells after 3 days of treatment of mixed culture is shown. Data are presented as mean ± s.d., n = 3.

Right panel: Relative survival of HMLE-Twist (purple, EMT) and HMLE-pBp (black, non-EMT) cells treated with salinomycin (Sal), GalNAc(α)-O-bn (GalNac-bn) or their combination at indicated concentrations. MTT assay was performed after 72 hours of treatment. Data are presented as mean ± s.d., n = 3 from quadruplicates. One-way ANOVA with post-hoc Tukey's multiple comparison test was used for statistical analysis (*$p < 0.05$, **$p < 0.01$, ***$p < 0.001$)

(E) Western blot analysis of membrane proteins N- and E-cadherin The effect of brefeldin A, thapsigargin and tunicamycin on N-cadherin and E-cadherin processing in EMT and non-EMT cells. Amidoblack staining of transferred proteins was used to evaluate equal loading.

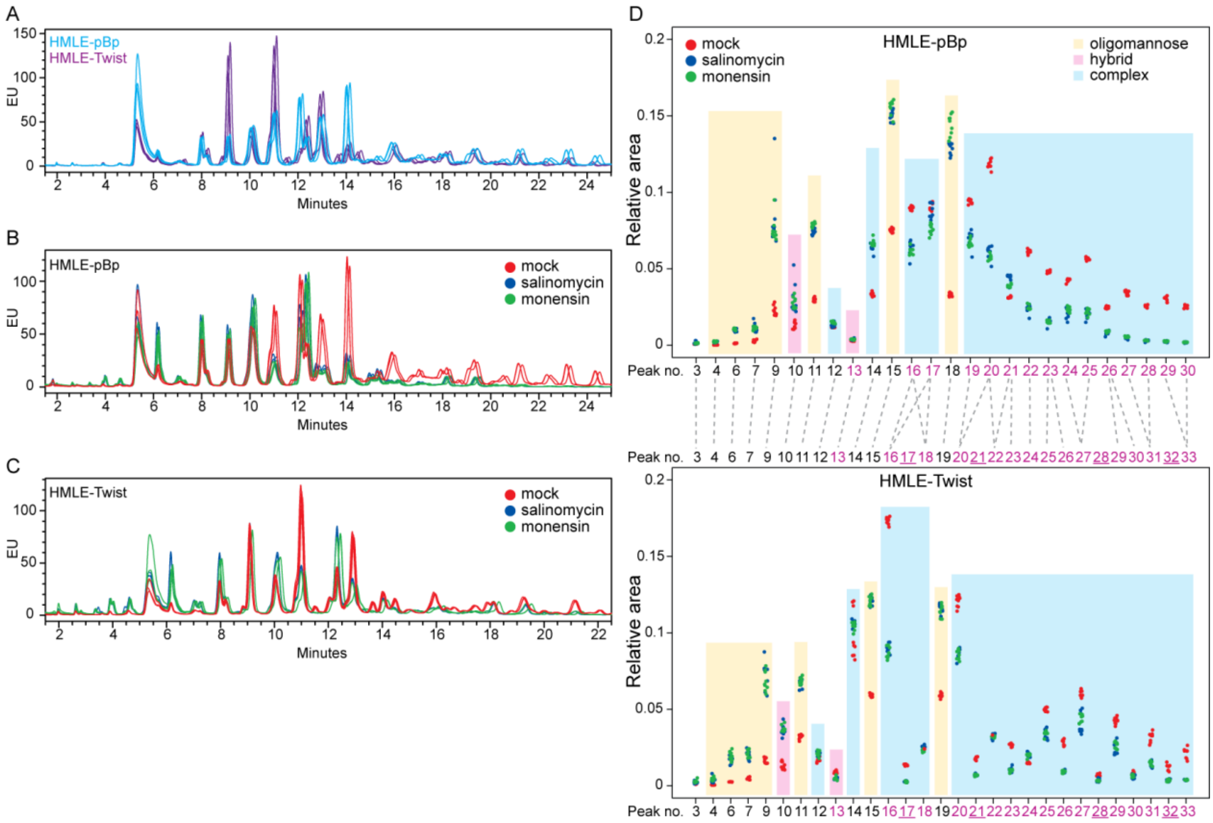


**Figure S5. Salinomycin affects N-glycosylation of secreted proteins.**
Overlaid chromatograms of HILIC-UPLC N-glycan profiles released from secreted proteins of mock treated EMT (HMLE-Twist) and non-EMT cells (HMLE-pBp) in (A); or mock, salinomycin and monensin treated non-EMT in (B) or EMT cells in (C). Triplicates of one representative experiment are shown in each panel.
Chromatograms in (A) were aligned by software to match the peak positions of EMT and non-EMT cells for straightforward comparison.
(D) Comparative analysis of N-glycans from secreted proteins of mock, salinomycin and monensin treated non-EMT (HMLE-pBp, upper panel) and EMT cells (HMLE-Twist, lower panel). Each area under the peak from chromatogram (B and C) was integrated and its relative amount plotted (n = 9; 3 technical replicates from each of 3 independent experiments). Peak numbers in magenta represent peaks containing sialylated glycans, underlined numbers denote peaks with sulfated/phosphorylated glycans (S/P), and grey dashed lines connect peaks containing matching major structures.

**Table S1. GSEA summary table with extended information**

Summary of GSEA analysis on different gene sets (ER and GA stressors or GO category) and their relation to enrichment in our experimental model (see Methods for details). * - significantly enriched; yellow - enrichment in mock treated sample, blue - enrichment in Sal treated sample. NES – normalized enrichment score, FDR – false discovery rate.

| **Cells** | **HMLE-pBp** | | | | **HMLE-Twist** | | | |
|---|---|---|---|---|---|---|---|---|
| **Experiment** | **CTRL vs. Sal** | | | | **CTRL vs. Sal** | | | |
| **Stressor or GO** | | *NES* | *FDR* | *p-value* | | *NES* | *FDR* | *p-value* |
| Tunicamycin | * | -1.42 | 0.107 | <0.01 | * | -1.35 | 0.093 | <0.01 |
| Thapsigargin | * | -1.38 | 0.083 | <0.01 | * | -1.24 | 0.117 | <0.01 |
| Doxorubicin | * | 1.28 | 0.24 | <0.01 | | 0.98 | 0.453 | 0.372 |
| 4MU-xyloside | * | -1.35 | 0.152 | 0.087 | | -0.97 | 0.526 | 0.569 |
| Brefeldin A | * | -1.3 | 0.132 | 0.17 | * | -1.43 | 0.06 | <0.01 |
| Monensin | | 0.82 | 0.762 | 0.611 | * | -1.2 | 0.124 | <0.01 |
| Nigericin | * | -1.37 | 0.083 | 0.143 | | -0.96 | 0.553 | 0.519 |
| HyT365 | * | -1.07 | 0.266 | 0.367 | * | -1.25 | 0.137 | <0.01 |
| Glycosylation | | 0.85 | 1 | 0.911 | * | -1.4 | 0.07 | <0.01 |

**Table S2**. **Differences in the relative amount of N-glycans of secreted proteins in non-EMT cells (HMLE-pBp).**

Glycan names are given in accordance with Oxford Glycobiology Institute notation: all N-glycans have core sugar sequence consisting of two N-Acetylglucosamines (GlcNAc) and three mannose residues. F in front of glycan name, indicates a core fucose α1–6 linked to the inner GlcNAc; Mx, number (x) of mannose on core GlcNAcs; Ax, number of antenna (GlcNAc) on trimannosyl core; B, bisecting GlcNAc linked β1–4 to β1–3 mannose; Gx, number of β1–4 linked galactose (G) on antenna; Sx, number (x) of N-Acetylneuraminic acids linked to galactose; S/P – sulfated/phosphorylated residue. Mean ± s.d. from n = 3 individual experiments is presented. Statistical significance was determined with one-way ANOVA with Tukey's multiple comparison post hoc test (ns – not significant, *p < 0.05, **p < 0.01, ***p < 0.001). Peaks containing only oligo-hexose sugars were eliminated from further analysis (peak numbers not present in Figure S5D and in Table below).

| Peak no. | Major N-glycan | Relative integrated area (%) | | | ANOVA | | |
|---|---|---|---|---|---|---|---|
| | | mock | salinomycin | monensin | mock vs sal | mock vs mon | sal vs mon |
| 1 | | n.d. | n.d. | n.d. | | | |
| 2 | | n.d. | n.d. | n.d. | | | |
| 3 | FM2 | 0,086 ± 0,001 | 0,153 ± 0,077 | 0,135 ± 0,014 | ns | ns | ns |
| 4 | M3 | 0,029 ± 0,004 | 0,212 ± 0,021 | 0,234 ± 0,033 | *** | *** | ns |
| 5 | | n.d. | n.d. | n.d. | | | |
| 6 | FM3 | 0,130 ± 0,014 | 0,977 ± 0,089 | 1,056 ± 0,049 | *** | *** | ns |
| 7 | M4 | 0,280 ± 0,014 | 1,107 ± 0,263 | 1,123 ± 0,072 | ** | ** | ns |
| 8 | | n.d. | n.d. | n.d. | | | |
| 9 | M5 | 2,289 ± 0,188 | 8,383 ± 1,444 | 7,549 ± 0,388 | ** | ** | ns |
| 10 | M4A1G1 | 1,292 ± 0,195 | 2,999 ± 0,819 | 2,793 ± 0,261 | * | ns | ns |
| 11 | M6 | 2,991 ± 0,100 | 7,389 ± 0,074 | 7,842 ± 0,131 | *** | *** | * |
| 12 | A2G2 | 1,200 ± 0,034 | 1,361 ± 0,083 | 1,505 ± 0,008 | * | ** | ns |
| 13 | M4A1G1S1 | 0,296 ± 0,012 | 0,376 ± 0,037 | 0,391 ± 0,020 | * | * | ns |
| 14 | FA2G2 | 3,337 ± 0,088 | 6,450 ± 0,217 | 6,702 ± 0,128 | *** | *** | ns |
| 15 | M7 | 7,562 ± 0,111 | 15,006 ± 0,229 | 15,570 ± 0,359 | *** | *** | ns |
| 16 | FA2G2S1 | 8,986 ± 0,059 | 6,374 ± 0,347 | 6,106 ± 0,118 | *** | *** | ns |
| 17 | FA2G2S1 | 8,991 ± 0,225 | 8,647 ± 0,546 | 7,588 ± 0,278 | ns | * | ns |
| 18 | M8 | 3,320 ± 0,084 | 12,883 ± 0,332 | 14,414 ± 0,609 | *** | *** | * |
| 19 | A2G2S2, FA2F1G2S1, FA2G2S2 | 9,405 ± 0,105 | 7,021 ± 0,369 | 6,665 ± 0,166 | *** | *** | ns |
| 20 | FA2G2S2 | 11,825 ± 0,237 | 6,104 ± 0,291 | 5,720 ± 0,038 | *** | *** | ns |
| 21 | FA2G2S2, FA3G3S1, A3G3S2 | 3,144 ± 0,053 | 4,416 ± 0,173 | 3,874 ± 0,050 | *** | ** | ** |
| 22 | A3G3S2, FA3G3S2 | 6,120 ± 0,062 | 2,417 ± 0,221 | 2,469 ± 0,109 | *** | *** | ns |
| 23 | FA3G3S2 | 4,822 ± 0,023 | 1,565 ± 0,154 | 1,510 ± 0,059 | *** | *** | ns |
| 24 | A3G3S3, FA3G3S3 | 4,261 ± 0,039 | 2,079 ± 0,287 | 2,335 ± 0,140 | *** | *** | ns |
| 25 | FA3G3S3, FA4G4S2, A3G3S3 | 5,663 ± 0,065 | 2,044 ± 0,242 | 2,188 ± 0,154 | *** | *** | ns |
| 26 | FA4G4S2 | 2,472 ± 0,057 | 0,814 ± 0,106 | 0,895 ± 0,050 | *** | *** | ns |
| 27 | FA4G4S3 | 3,427 ± 0,121 | 0,499 ± 0,059 | 0,559 ± 0,030 | *** | *** | ns |
| 28 | FA4G4S3 | 2,534 ± 0,058 | 0,296 ± 0,033 | 0,315 ± 0,004 | *** | *** | ns |
| 29 | FA4G4S4 | 3,033 ± 0,138 | 0,238 ± 0,028 | 0,264 ± 0,025 | *** | *** | ns |
| 30 | FA4G4S4 | 2,540 ± 0,059 | 0,188 ± 0,011 | 0,198 ± 0,009 | *** | *** | ns |

**Table S3**. **Differences in the relative amount of N-glycans of secreted proteins in EMT cells (HMLE-Twist).**

Glycan names are given in accordance with Oxford Glycobiology Institute notation: all N-glycans have core sugar sequence consisting of two N-Acetylglucosamines (GlcNAc) and three mannose residues. F in front of glycan name, indicates a core fucose α1–6 linked to the inner GlcNAc; Mx, number (x) of mannose on core GlcNAcs; Ax, number of antenna (GlcNAc) on trimannosyl core; B, bisecting GlcNAc linked β1–4 to β1–3 mannose; Gx, number of β1–4 linked galactose (G) on antenna; Sx, number (x) of N-Acetylneuraminic acids linked to galactose; S/P – sulfated/phosphorylated residue. Mean ± s.d. from n = 3 individual experiments is presented. Statistical significance was determined with one-way ANOVA with Tukey's multiple comparison post hoc test (ns – not significant, *p < 0.05, **p < 0.01, ***p < 0.001). Peaks containing only oligo-hexose sugars were eliminated from further analysis (peak numbers not present in Figure S5D and in Table below).

| Peak no. | Major N-glycan | Relative integrated area (%; average ± s.d.) | | | ANOVA | | |
|---|---|---|---|---|---|---|---|
| | | mock | salinomycin | monensin | mock vs sal | mock vs mon | sal vs mon |
| 1 | | n.d. | n.d. | n.d. | | | |
| 2 | | n.d. | n.d. | n.d. | | | |
| 3 | FM2 | 0,117 ± 0,009 | 0,225 ± 0,045 | 0,288 ± 0,060 | ns | * | ns |
| 4 | M3 | 0,050 ± 0,005 | 0,374 ± 0,123 | 0,427 ± 0,079 | * | ** | ns |
| 5 | | n.d. | n.d. | n.d. | | | |
| 6 | FM3 | 0,256 ± 0,003 | 1,708 ± 0,198 | 1,960 ± 0,245 | *** | *** | ns |
| 7 | M4, FA1 | 0,477 ± 0,067 | 1,994 ± 0,255 | 2,076 ± 0,203 | *** | *** | ns |
| 8 | | n.d. | n.d. | n.d. | | | |
| 9 | M5 | 1,642 ± 0,157 | 7,305 ± 0,738 | 6,835 ± 0,370 | *** | *** | ns |
| 10 | M4A1G1 | 1,315 ± 0,227 | 3,653 ± 0,271 | 3,735 ± 0,074 | *** | *** | ns |
| 11 | M6 | 3,195 ± 0,065 | 6,731 ± 0,250 | 6,849 ± 0,145 | *** | *** | ns |
| 12 | A2G2 | 1,719 ± 0,098 | 2,154 ± 0,128 | 2,127 ± 0,125 | * | * | ns |
| 13 | M4A1G1S1 | 0,901 ± 0,034 | 0,458 ± 0,024 | 0,479 ± 0,031 | *** | *** | ns |
| 14 | FA2G2 | 9,864 ± 1,512 | 10,632 ± 0,122 | 10,443 ± 0,162 | ns | ns | ns |
| 15 | M7 | 5,902 ± 0,082 | 12,103 ± 0,285 | 12,048 ± 0,246 | *** | *** | ns |
| 16 | FA2G2S1 | 17,329 ± 0,072 | 8,956 ± 0,238 | 8,701 ± 0,322 | *** | *** | ns |
| 17 | FA2G2S1(S/P) | 1,357 ± 0,037 | 0,277 ± 0,028 | 0,261 ± 0,014 | *** | *** | ns |
| 18 | FA2G2S1 | 2,383 ± 0,022 | 2,542 ± 0,027 | 2,320 ± 0,054 | * | ns | ** |
| 19 | M8 | 5,933 ± 0,107 | 11,543 ± 0,270 | 11,475 ± 0,228 | *** | *** | ns |
| 20 | FA2G2S2 | 12,072 ± 0,325 | 8,599 ± 0,218 | 8,561 ± 0,262 | *** | *** | ns |
| 21 | FA2G2S2(S/P) | 1,799 ± 0,087 | 0,721 ± 0,029 | 0,696 ± 0,013 | *** | *** | ns |
| 22 | FA3G3S1 | 3,309 ± 0,017 | 3,242 ± 0,079 | 3,201 ± 0,034 | ns | ns | ns |
| 23 | FA3G3S1 | 2,670 ± 0,100 | 1,068 ± 0,059 | 0,965 ± 0,040 | *** | *** | ns |
| 24 | FA3G3S1, A3G3S2 | 1,499 ± 0,041 | 2,054 ± 0,103 | 1,993 ± 0,025 | *** | *** | ns |
| 25 | FA3G3S2 | 5,000 ± 0,096 | 3,322 ± 0,309 | 3,447 ± 0,096 | *** | *** | ns |
| 26 | FA3G3S2 | 2,857 ± 0,167 | 0,923 ± 0,067 | 0,921 ± 0,024 | *** | *** | ns |
| 27 | FA4G4S1, A3G3S3, FA3G3S3 | 5,989 ± 0,173 | 4,030 ± 0,625 | 4,398 ± 0,201 | ** | * | ns |
| 28 | FA3G3S3(S/P) | 0,713 ± 0,057 | 0,343 ± 0,081 | 0,326 ± 0,008 | ** | ** | ns |
| 29 | A3G3S3, FA4G4S2 | 4,257 ± 0,193 | 2,363 ± 0,361 | 2,634 ± 0,180 | *** | ** | ns |
| 30 | FA4G4S2 | 0,847 ± 0,054 | 0,557 ± 0,116 | 0,637 ± 0,057 | * | ns | ns |
| 31 | FA4G4S3 | 3,187 ± 0,269 | 1,368 ± 0,169 | 1,452 ± 0,105 | *** | *** | ns |
| 32 | FA4G4S3(S/P) | 1,217 ± 0,175 | 0,367 ± 0,043 | 0,352 ± 0,037 | *** | *** | ns |
| 33 | FA4G4S4 | 2,146 ± 0,291 | 0,389 ± 0,015 | 0,394 ± 0,006 | *** | *** | ns |

**Additional file S1 (separate file). List of selective compounds from the initial screen**
(A) List of 227 EMT selective compounds from screen by Gupta et al. obtained by our modified algorithm.
(B) Selectivity of monensin and lasalocid A compared to published compounds from screen by Gupta et al. (6).

**Additional file S2 (separate file). Differential expression and gene ontology analysis of RNA-seq data**
(A – C) Complete list of differentially expressed genes in EMT and non-EMT cells with or without treatment with Sal. List of genes differentially expressed between un-treated HMLE-pBp and HMLE-Twist cells (A), HMLE-pBp cells treated with 0.2 μM Sal (B) or HMLE-Twist cells treated with 0.2 μM Sal (C).
(D and E) Analysis of ≥2 fold change differentially expressed genes in EMT and non-EMT cells after treatment with Sal.
(D) List of genes that are differentially expressed (≥ 2 fold change) in RNA-seq experiment after treatment with Sal. List is divided by Venn diagram showing overlapping and non-overlapping genes between experiments (see Figure S2A).
(E) List of 64 differentially expressed genes after Sal (≥ 2 fold change UP in EMT & unchanged or DOWN in non-EMT cells). See Figure S2B and text for more details.
(F) PANTHER overrepresentation test for biological process and cellular component GO categories on the 52 gene set (2-fold change UP in both EMT and non-EMT cells after Sal).
(G) PANTHER overrepresentation test for biological process and cellular component GO categories on the 52 gene set (≥ 2 fold change UP in EMT and non-EMT cells after Sal).
(H – J) List of biological process GO categories in EMT and non-EMT cells after treatment with Sal.
(H) List of GO categories overlapping between experimental setup's (EMT vs non-EMT, EMT vs Sal, non-EMT vs Sal).
(I) GO categories unique for Sal treated EMT cells sorted by p-value.
(J) GO categories unique for Sal treated non-EMT cells sorted by p-value.

**Additional file S3 (separate file). Detected N-glycan structures**
Compositions and proposed structures of N-glycans of the secreted proteins from mock, Sal and Mon (0.2 μM) treated EMT (A) and non-EMT (B) cells. Multiple N-glycan structures found within the same peak were divided into major structure (most abundant), lower abundance (~below 50% relative to the most abundant structure) and low abundance (~below 20% relative to the most abundant structure). Composition labels: N – GlcNAc, H – mannose or galactose, F – fucose, S – sialic acid and S/P – sulfated/phosphorylated residue.
